# *NvPrdm14d*-expressing neural progenitor cells contribute to non-ectodermal neurogenesis in *Nematostella vectensis*

**DOI:** 10.1101/2022.07.06.498948

**Authors:** Quentin I. B. Lemaître, Natascha Bartsch, Ian U. Kouzel, Henriette Busengdal, Gemma Sian Richards, Patrick R. H. Steinmetz, Fabian Rentzsch

**Affiliations:** Sars International Centre for Marine Molecular Biology, University of Bergen, Thormøhlensgate 55, 5006 Bergen, Norway; Department for Biological Sciences, University of Bergen, Thormøhlensgate 55, 5006 Bergen, Norway

**Keywords:** Cnidaria, PR domain, endodermal neurogenesis, mesodermal neurogenesis, non- ectodermal neurogenesis, motoneuron, motor neuron

## Abstract

Neurogenesis has been studied extensively in the ectoderm, from which most animals generate the majority of their neurons. Neurogenesis from non-ectodermal tissue is, in contrast, poorly understood. Here we use the cnidarian *Nematostella vectensis* as a model to provide new insights into the molecular regulation of non-ectodermal neurogenesis. We show that the transcription factor *NvPrdm14d* is expressed in a subpopulation of *NvSoxB(2)-*expressing endodermal progenitor cells and their *NvPOU4*-expressing progeny. Using a new transgenic reporter line, we show that *NvPrdm14d*-expressing cells give rise to neurons in the body wall and in close vicinity of the longitudinal retractor muscles. RNA-sequencing of *NvPrdm14d*::GFP-expressing cells and gene knockdown experiments provide candidate genes for the development and function of these neurons. Together, the identification of a population of endoderm-specific neural progenitor cells and of previously undescribed putative motoneurons in *Nematostella* provide new insights into the regulation of non-ectodermal neurogenesis.

## INTRODUCTION

Adult animals contain neural cells located in tissues derived from all embryonic germ layers whereas neurogenesis, the generation of these neural cells, occurs in most animals almost exclusively in the ectoderm. While non-ectodermal neurogenesis occurs in several taxa (**Figure 1A**), it typically contributes only a small fraction of neurons. Even the non-ectodermal (*i.e.* mesodermal and endodermal) aspects of the nervous system mainly consist of neurons generated by neural progenitors or precursors of ectodermal origin that migrated into these germ layers (Bronner-Fraser and Fraser, 1991; Mayor and Theveneau, 2012; Nagy and Goldstein, 2017). The molecular regulation of non-ectodermal neurogenesis has been studied in only a few cases and it remains therefore unclear to what extent ectodermal and non-ectodermal neurogenesis differ within a given species, and whether conserved regulators of non-ectodermal neurogenesis exist (Henrique et al., 2015; Luo and Horvitz, 2017; Richards and Rentzsch, 2015; Wei et al., 2011).

**Figure 1:**
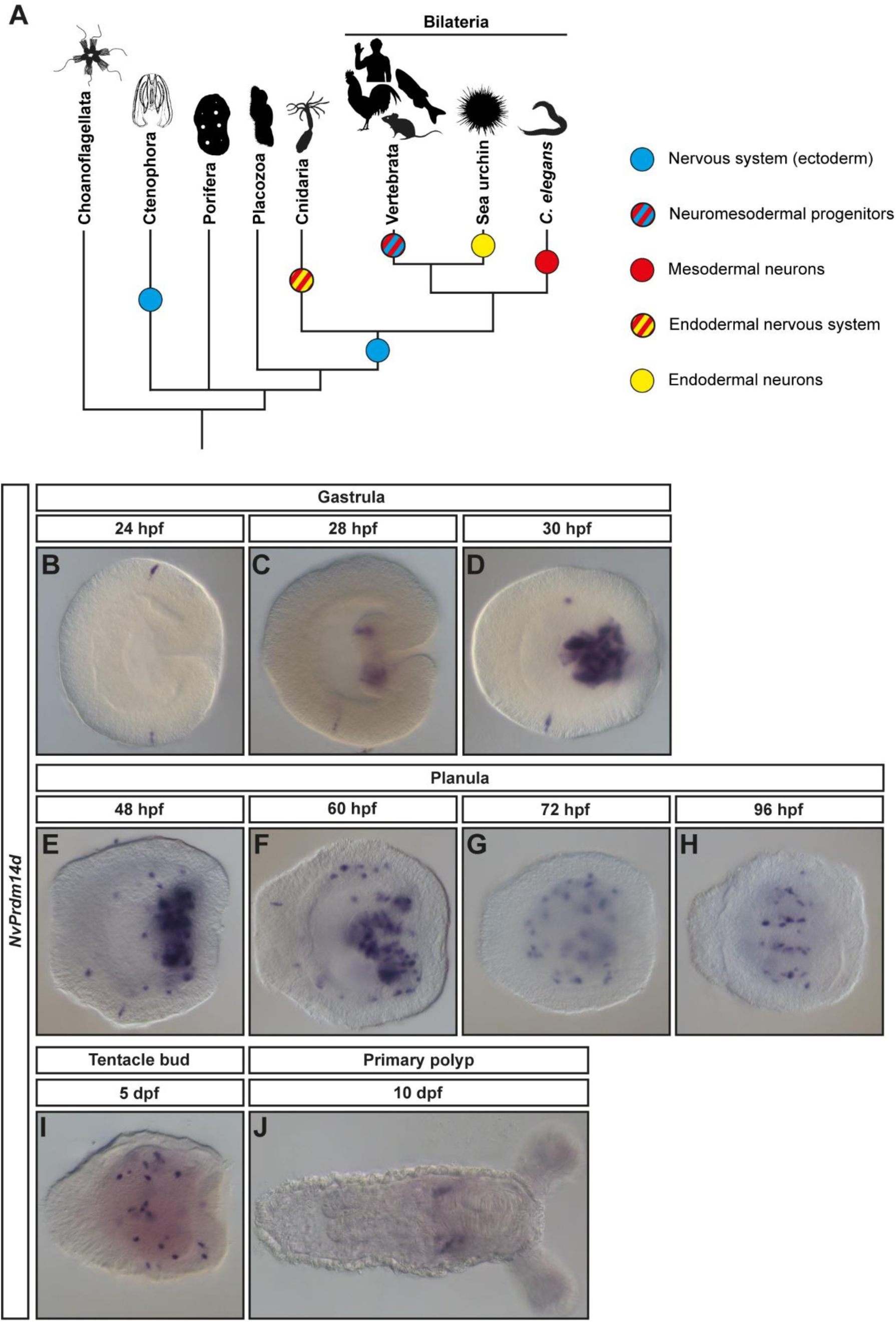
NvPrdm14d is mainly expressed in scattered endodermal cells. (**A**) Simplified phylogeny showing the distribution of ectodermal and non-ectodermal neurogenesis across metazoans. Presence of a nervous system is indicated by a blue dot, independently from a single or multiple origin(s) of the nervous system. Non-ectodermal neurogenesis is indicated with dots of various colors illustrating the germ layer producing non-ectodermal neurons. Except for hydrozoan cnidarians, all cases of non- ectodermal neurogenesis are occurring in addition to canonical ectodermal neurogenesis. When applicable, cartoons show representative species in which non-ectodermal neurogenesis have been studied. The phylogeny is rooted with choanoflagellates as outgroup, and is based on (Dunn et al., 2014). Animal silhouettes are from https://phylopic.org/. (**B-J**) Expression pattern of NvPrdm14d by colorimetric in situ hybridization. (**B**) In the early gastrula, NvPrdm14d is expressed in very few single ectodermal cells. (**C, D**) In mid-gastrula, NvPrdm14d starts to be expressed in the pharynx. Strong expression in the pharynx persists until mid-planula (**F**). In early planula (**E**), NvPrdm14d starts to be expressed in scattered endodermal cells. (**G-I**) From mid-planula, the expression remains in scattered cells within the pharynx and the endoderm. (**J**) In the primary polyp, NvPrdm14d is expressed in lateral domains within the endoderm. In all pictures, the oral pole is oriented to the right.

In vertebrates, neuromesodermal progenitors (NMps) are bipotent cells located in the tailbud and they are involved in axial elongation (Henrique et al., 2015). These cells are presumably derived from mesodermal tissues and have the potential to produce both the posterior spinal cord and paraxial mesoderm (Tzouanacou et al., 2009). The bipotency of these NMps is maintained by the co-expression of the neural marker gene *Sox2* and the early mesodermal marker gene *T/Brachyury* (Garriock et al., 2015; Olivera-Martinez et al., 2012; Tsakiridis et al., 2014). NMps are specified to either a neural or paraxial mesoderm fate depending on the extrinsic cue they receive when leaving their niche. Anteriorly, cells are exposed to high levels of retinoic acid and initiate neural differentiation, while posteriorly, they are exposed to high levels of Wnt3a and initiate mesodermal differentiation (Cambray and Wilson, 2002; Cunningham et al., 2015; Garriock et al., 2015; Niederreither et al., 2000; Selleck and Stern, 1991; Takada et al., 1994; Wymeersch et al., 2016; Yoshikawa et al., 1997). These NMps contribute to posterior neural structures and to the posterior somites (Albors et al., 2018; Cunningham et al., 2015; Garriock et al., 2015; Jurberg et al., 2014).

In the nematode *C. elegans*, 6 out of the 20 pharyngeal neurons are derived from the mesodermal lineage MS, including the I4 neurons (Sulston et al., 1983). It has been shown that the I4 neurons are specified in part through the action of the proneural protein HLH-3 and the HLH-2/Mediator complex repressing the CDK-7/Cyclin-H complex (Luo and Horvitz, 2017). However, the specification of the other mesodermal neurons remains to be investigated.

In the sea urchin, the endodermal origin of the foregut neurons has been demonstrated by using photoconvertible fluorescent proteins (Wei et al., 2011). In the endoderm, the expression of *SoxB1* maintains the neurogenic potential and induces the expression of *Six3*. Then, *Six3* activates the *SoxC*-mediated and *Nkx3-2*-mediated neurogenic cascades. While the *SoxC* cascade is used in both ectodermal and endodermal neurogenic pathways, the *Nkx3-2* cascade seems to be specific to the endodermal neurogenic pathway. It is also apparent that Nodal, BMP, Notch, FGF and Wnt signaling are involved in sea urchin endodermal neurogenesis but further investigations are required to understand their respective interactions within the endodermal neurogenic pathway (McClay et al., 2018; Wei et al., 2009; Wei et al., 2011; Wei et al., 2016).

Cnidarians, the sister group to bilaterians, develop from only two germ layers, here called ectoderm and endoderm. The cnidarian endoderm has been suggested to be homologous to either both endoderm and mesoderm, or only to the mesoderm of bilaterians (Martindale et al., 2004; Steinmetz et al., 2017; Technau, 2020). The nervous system of cnidarian polyps lacks brain-like centralization and consists of sensory/sensory-motor neurons, ganglion neurons (morphologically similar to interneurons) and cnidocytes, the phylum-specific stinging cells (Galliot et al., 2009; Rentzsch et al., 2016; Watanabe et al., 2009). Non-ectodermal neurogenesis is a common feature in cnidarians. Hydrozoan cnidarians, such as *Hydra* and *Hydractinia*, generate the entirety of their nervous system from an endoderm-derived cell type: the interstitial cells (Genikhovich et al., 2006; Gröger and Schmid, 2001; Leclère et al., 2012; Martin and Archer, 1986; Martin and Thomas, 1981; Plickert et al., 1988; Weis et al., 1985). In anthozoan cnidarians, notably the sea anemone *Nematostella vectensis*, the nervous system is found in both the ectoderm and the endoderm (Marlow et al., 2009; Nakanishi et al., 2012; Richards and Rentzsch, 2014). In *Nematostella*, the endodermal origin of non-ectodermal neurons has been demonstrated by transplantation experiments (Nakanishi et al., 2012). Perturbation experiments showed that Notch signaling and the transcription factors *NvAth-like* and *NvSoxB(2)* act at an early stage of both ectodermal and endodermal neurogenesis, with Notch signaling acting as a negative and *NvAth-like* and *NvSoxB(2)* as positive regulators, respectively (Layden and Martindale, 2014; Richards and Rentzsch, 2014; Richards and Rentzsch, 2015). Downstream of these factors, *NvAshA*, *NvDmrtB* and *NvPOU4* were shown to be expressed and function in large populations of ectodermal and endodermal neural cells (Layden et al., 2012; Parlier et al., 2013; Tournière et al., 2020). However, the molecular mechanisms underlying more specifically endodermal neurogenesis in *Nematostella* have not been investigated so far. In the present study, we show that *NvPrdm14d* is involved in the generation of a subpopulation of endodermal neurons.

Prdm14 belongs to the PR domain (<u>P</u>RDI-BF1 and <u>R</u>IZ1 homology domain) containing family of transcription factors (Buyse et al., 1995; Hohenauer and Moore, 2012; Keller and Maniatis, 1991). Prdm proteins are composed of a PR domain related to the catalytic SET domain (<u>S</u>uppressor of variegation 3-9, <u>E</u>nhancer of zeste and <u>T</u>rithorax) characterizing a large group of histone lysine methyltransferases (HMT), followed by a variable number of zinc fingers at the C-terminus (Fumasoni et al., 2007; Hohenauer and Moore, 2012; Huang, 2002; Kinameri et al., 2008; Sun et al., 2008; Wu et al., 2010). While the PR domain of Prdm14 does not show intrinsic HMT activity, it recruits co-factors, whereas the zinc fingers directly bind to regulatory elements of target genes (Chia et al., 2010; Liu et al., 2012; Ma et al., 2011; Yamaji et al., 2013).

In vertebrates, Prdm14 is a major factor involved in the maintenance of embryonic stem cells. It recruits TET (<u>T</u>en-<u>E</u>leven <u>T</u>ranslocation protein) to the promoter of pluripotency genes (*e.g. Pou5f1/Oct4*, *Nanog*, *Sox2*, *Klf5*) for promoting their expression, while in parallel, it recruits PRC2 (<u>P</u>olycomb <u>R</u>epressive <u>C</u>omplex 2) for repressing differentiation genes (Chia et al., 2010; Ma et al., 2011; Nakaki and Saitou, 2014; Okashita et al., 2014; Tsuneyoshi et al., 2008; Wu and Zhang, 2014; Yamaji et al., 2013). Furthermore, Prdm14 can function in the reprogramming of differentiated cells to induce pluripotent stem cells (iPS) *in vitro*. (Chia et al., 2010; Geijsen, 2012; Gillich et al., 2012; Nakaki et al., 2013; Takahashi et al., 2007).

Similarly, Prdm14 is required for the specification of primordial germ cells (PGCs) in vertebrates as it allows the reacquisition of pluripotency. It acts in synergy with Prdm1 and TFAP2C (encoding AP2γ) to promote pluripotency and germline-specific gene expression, while repressing somatic genes (Magnúsdóttir et al., 2013; Nakaki et al., 2013; Sybirna et al., 2020; Yamaji et al., 2008). The absence of Prdm14 leads to a deficiency of PGCs and therefore, results in sterility (Hadziselimovic et al., 2018; Ohinata et al., 2009; Okuzaki et al., 2019; Sybirna et al., 2020; Yamaji et al., 2008).

In the zebrafish, Prdm14 is involved in axon outgrowth of primary motoneurons through the activation of *Islet2* (Liu et al., 2012). A role for *Prdm14* and *Islet2* in the development of motoneurons has also been described in amphioxus, a non-vertebrate chordate (Kawaguchi et al., 2019). These observations led to the suggestion that the ancestral function of *Prdm14* is in neural development and that it has been co-opted for the regulation of pluripotent cells during vertebrate evolution (Kawaguchi et al., 2019).

In the present report, we identify *NvPrdm14d* as a factor involved in endodermal neurogenesis in *Nematostella*. By combining colorimetric and fluorescent *in situ* hybridization with EdU labelling, we show that *NvPrdm14d* is expressed in a subpopulation of non-ectodermal neural progenitor cells (NPCs), as well as in the progeny of these cells. We generated a *NvPrdm14d::GFP* transgenic reporter line, which highlights a population of putative motoneurons in the close vicinity of retractor muscles. We then sorted *NvPrdm14d*::GFP-expressing cells and generated their transcriptome to identify additional potential candidate genes involved in the generation of such neurons and tested the requirement of *NvPrdm14d* for their expression by shRNA injection.

## RESULTS

### *NvPrdm14d* is a candidate gene involved in endodermal neurogenesis

The genome of *Nematostella vectensis* contains four well conserved *Prdm14* paralogs [*NvPrdm14a-d,* (Vervoort et al., 2015)]. Inspection of diverse transcriptome datasets revealed that each *NvPrdm14* paralog is overrepresented in at least one neural transcriptome (Error! Reference source not found.), but by *in situ* hybridization (ISH), we obtained a clear expression pattern only for *NvPrdm14d*.

We found that *NvPrdm14d* is expressed in very few single ectodermal cells at early gastrula stage (**Figure 1B**), before starting to be expressed in individual cells in the pharynx from mid-gastrula (**Figure 1C**). The colorimetric *in situ* hybridizations did not allow to decide unambiguously whether the pharyngeal staining is located in the ectodermal, the endodermal or both parts of the pharynx. The strong expression in cells in the pharynx lasts until the mid-planula stage (**Figure 1F**). From early planula stage, *NvPrdm14d* is expressed in scattered endodermal cells in the body wall (**Figure 1E**) and from mid-planula, it is expressed in these scattered endodermal and some pharyngeal cells (**Figure 1G-I**). In addition, *NvPrdm14d* is expressed in a small number of ectodermal cells during gastrula and planula stages (**Figure 1C- I**). In the primary polyp, *NvPrdm14d* is expressed in some endodermal domains on the oral side (**Figure 1J**). Overall, the expression pattern in scattered cells and the onset of endodermal *NvPrdm14d*-expression at the stage when the first neural progenitor cells (NPCs) appear in the *Nematostella* endoderm (Richards and Rentzsch, 2014) are consistent with a role for *NvPrdm14d* in endodermal neurogenesis.

### *NvPrdm14d* is expressed in a subset of mesendodermal NPCs and their progeny

To characterize the expression pattern of *NvPrdm14d* in more detail, we performed double fluorescence *in situ* hybridization (DFISH). The higher resolution afforded by DFISH and confocal imaging showed that the pharyngeal expression of *NvPrdm14d* is almost exclusively observed in the endodermal part of the pharynx (**Figure 2A, E, I** and **M**). We then compared the expression pattern of *NvPrdm14d* with *NvSoxB(2)*, a gene that is expressed in a population of progenitor cells that give rise to neurons and gland/secretory cells (neural/secretory progenitor cells, N/SPCs) in both the ectoderm and endoderm in *Nematostella* (Richards and Rentzsch, 2014; Steger et al., 2022; Tournière et al., 2022). At planula stage, quantification of confocal images showed that the number of cells expressing *NvSoxB(2)* was higher than that expressing *NvPrdm14d* (**Figure 2A**). While on average half of the *NvPrdm14d*- expressing cells also express *NvSoxB(2),* only a minority of *NvSoxB(2)*-expressing cells in the endoderm co-express *NvPrdm14d* (**Figure 2A-D**). This suggests that *NvPrdm14d* is expressed in a subset of endodermal progenitor cells.

**Figure 2:**
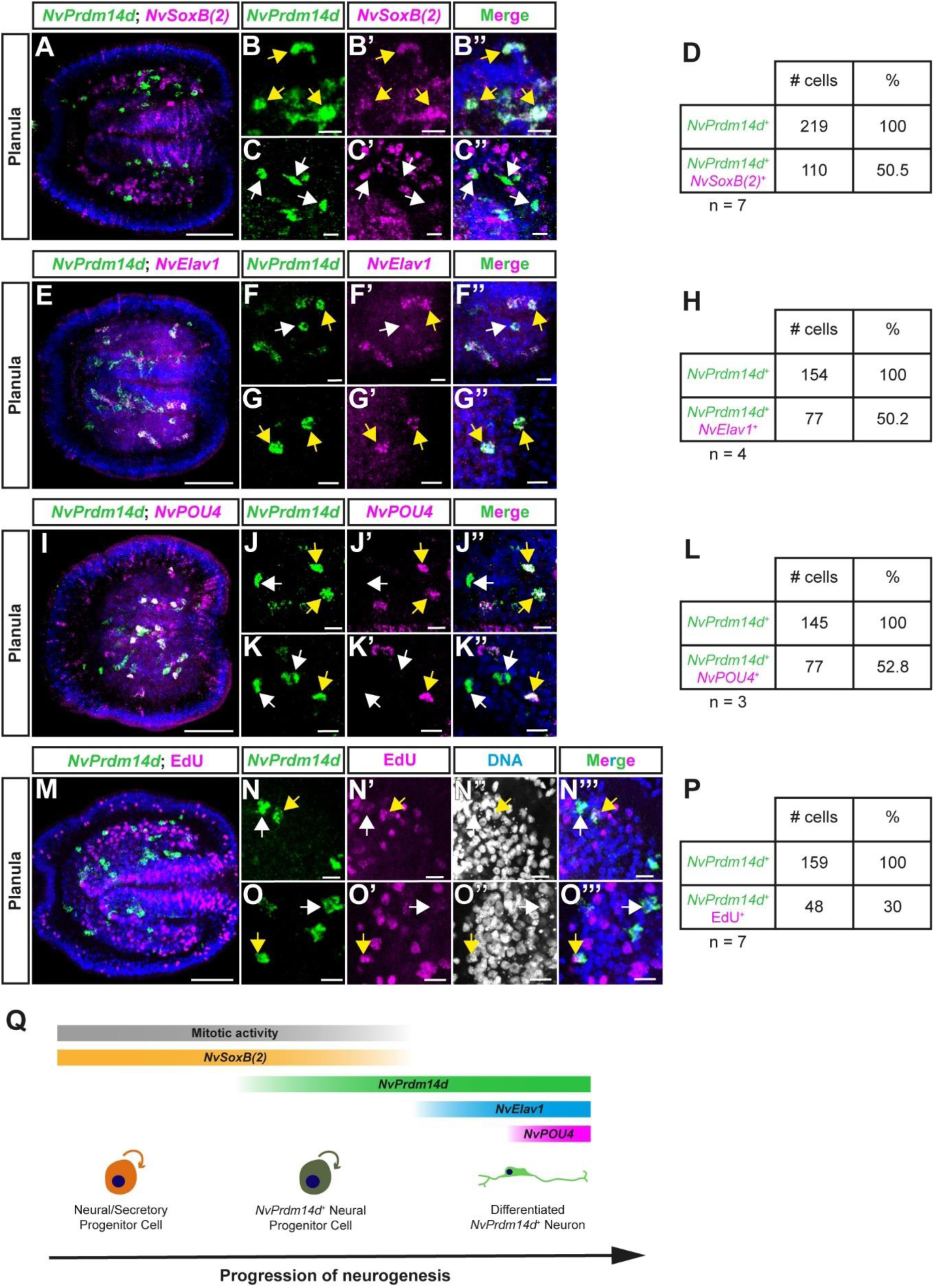
NvPrdm14d is expressed in a subset of endodermal neural/secretory progenitor cells as well as in their progeny. (**A-C**, **E-G & I-K**) Confocal images of double fluorescence in situ hybridization for NvPrdm14d (green) and different neural marker genes (magenta). (**B-C**) Enlargements of NvPrdm14d^+^ cells co-expressing (or not) NvSoxB(2). (**D**) Quantification of NvPrdm14d^+^ cells co-expressing NvSoxB(2): 15-76% (average 50.5%, n=7) of NvPrdm14d^+^ cells co-express NvSoxB(2). (**F-G**) Enlargements of NvPrdm14d^+^ cells co-expressing (or not) NvElav1. (**H**) Quantification of NvPrdm14d^+^ cells co-expressing NvElav1: 41-56% (average 50.2%, n=4) of NvPrdm14d^+^ cells co-express NvElav1. (**J-K**) Enlargements of NvPrdm14d^+^ cells co-expressing (or not) NvPOU4. (**L**) Quantification of NvPrdm14d^+^ cells co-expressing NvPOU4: 50%-55% (average 52.8%, n=3) of NvPrdm14d^+^ cells co-express NvPOU4. (**M-O**) Confocal images of fluorescence in situ hybridization for NvPrdm14d combined with the fluorescent labelling of proliferating cells by nuclear EdU staining, respectively shown in green and magenta. (**N-O**) Enlargements showing that only few NvPrdm14d^+^ cells are co-labelled by EdU. (**P**) Quantification of NvPrdm14d expressing cells co-labeled by EdU: 21-42% (average 30%, n=7) of NvPrdm14d^+^ cells are EdU positive. (**Q**) Schematics summarizing the temporal expression of NvPrdm14d. The expression starts in a subset of endodermal neural/secretory progenitor cells that are still dividing and expressing NvSoxB(2). The expression of NvPrdm14d continues in neurons expressing differentiation markers such as NvElav1 and NvPOU4. Yellow arrows indicate co-labelled cells, while white arrows indicate cells with a single label. Embryos are counterstained for DNA in blue, the oral pole is oriented to the right, scale bars: 50 µm for full embryos (**A**, **E**, **I & M**), 10 µm for enlargements (**B-C**, **F-G**, **J-K & N-O**).

Additionally, we compared the expression pattern of *NvPrdm14d* with the neuronal differentiation markers *NvElav1* and *NvPOU4* (Nakanishi et al., 2012; Tournière et al., 2020) to determine whether the expression of endodermal *NvPrdm14d* is restricted to progenitor cells or lasts until neurons differentiate. At planula stage, we found that on average about 50% of the *NvPrdm14d*-expressing cells co-express *NvElav1* (**Figure 2E-H**). We observed a similar proportion of *NvPrdm14d^+^* cells expressing *NvPOU4* (**Figure 2I-L**), consistent with the co-expression of *NvElav1* and *NvPOU4* in endodermal neurons (Tournière et al., 2020). This indicates that the expression of *NvPrdm14d* persists in differentiated endodermal neurons derived from the *NvPrdm14d*^+^ endodermal NPCs.

To test whether *NvPrdm14d* is expressed in proliferative cells, we combined FISH with EdU labelling. It has been previously determined that the time from S-phase until mitosis of *NvSoxB(2)^+^* N/SPCs is approximately 4 hours (Richards and Rentzsch, 2014). Therefore, we incubated planula larvae in EdU for 4 hours to label most of the potentially dividing *NvPrdm14d^+^* cells. At planula stage, we found that on average 30% of the *NvPrdm14d^+^* cells are EdU positive (**Figure 2M-P**), showing that at least some *NvPrdm14d^+^* cells are dividing.

Altogether, our data suggest that the expression of *NvPrdm14d* starts in a subset of dividing *NvSoxB(2)*^+^ endodermal N/SPCs. Its expression persists in differentiated endodermal neurons after the expression of *NvSoxB(2)* ceased (**Figure 2Q**).

### *NvPrdm14d*^+^ cells differentiate mainly into endodermal ganglion neurons

To allow the visualization of the morphology of *NvPrdm14d*^+^ N/SPCs and their progeny, we generated a transgenic reporter line (Renfer and Technau, 2017; Renfer et al., 2010) consisting of a gene encoding a membrane-tethered GFP under the control of a 5 kb sequence from the regulatory region of *NvPrdm14d,* referred to as *NvPrdm14d::GFP*.

DFISH for *NvPrdm14d* and GFP mRNAs in *NvPrdm14d::GFP* transgenic planulae showed that most *NvPrdm14d*^+^ cells co-express GFP mRNA and that all cells expressing GFP mRNA are *NvPrdm14d*^+^ (**Figure 3A-C**). We found, however, that some *NvPrdm14d*^+^ cells do not co-express GFP mRNA, particularly in the pharynx (**Figure 3A-C**).

**Figure 3:**
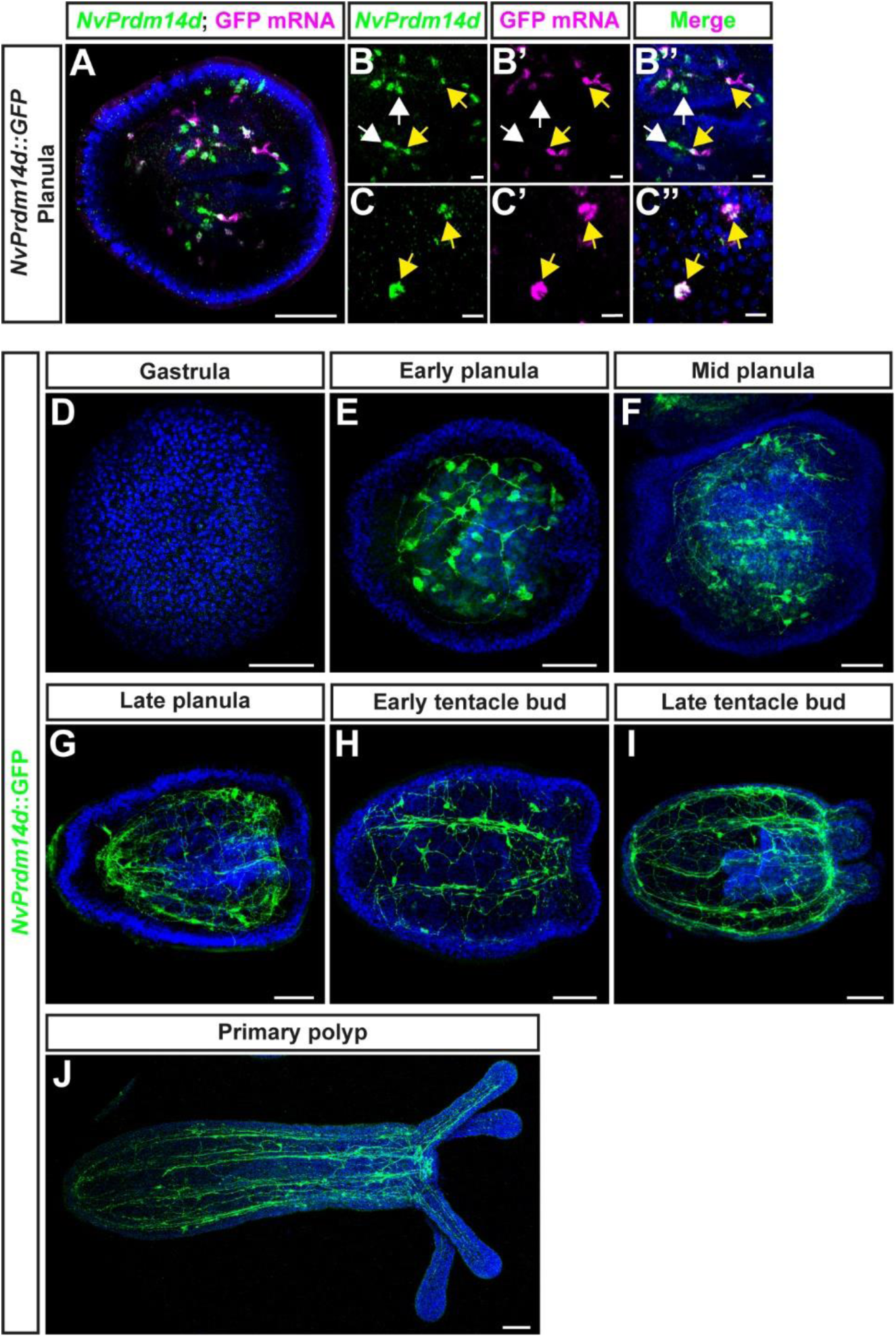
A transgenic reporter line for NvPrdm14d highlights a population of endodermal neurons. (**A-C**) Confocal images of double fluorescence in situ hybridization for NvPrdm14d (green) and GFP mRNA (magenta) in NvPrdm14d::GFP transgenic embryos. (**B-C**) Enlargements of NvPrdm14d^+^ cells co- expressing (or not) GFP mRNA. Most GFP^+^ cells co-express NvPrdm14d (yellow arrows). However, several NvPrdm14d^+^ cells are GFP^-^, notably in the pharynx (white arrows). (**D-J**) Time course of confocal images of immunofluorescence staining for NvPrdm14d::GFP over developmental stages revealing neurons forming a subset of the endodermal nerve net and of the neurites composing the longitudinal tracts. GFP is shown in green. Samples are counterstained for DNA in blue. In all pictures, the oral pole is oriented to the right. Scale bars: 50 µm for full embryos (**A & D-H**), 10 µm for enlargements (**B-C**).

The analysis of *NvPrdm14d*::GFP expressing animals identified GFP^+^ neurons in the endoderm (**Figure 3D-J**). Expression of the transgene can be visualized from early planula in cells exhibiting neurite projections, however some GFP^+^ cells lack neurites at this stage (**Figure 3E & 4A**). During larval development, an increasing number of neurons are labelled by GFP, and neurites progressively form longitudinal tracts along the mesenteries as well as the endodermal nerve net (**Figure 3F-I**). Additionally, we detect GFP^+^ neurites in the tentacles of the primary polyp (**Figure 3J**). Compared to previously described transgenic reporter lines for *NvSoxB(2)* and *NvElav1* (Nakanishi et al., 2012; Richards and Rentzsch, 2014), the *NvPrdm14d* reporter labels a smaller number of cells. We did not observe cells with the morphology of gland/secretory cells.

Next, we looked more closely at individual GFP^+^ cells, and we noticed that some of the cells lacking projections in early planulae, are dividing (**Figure 4A-A’’**). We crossed the *NvPrdm14d::GFP* line with the previously described *NvSoxB(2)::mOrange* line to obtain double transgenics (Richards and Rentzsch, 2014). In such animals, the GFP^+^ dividing cells are also mOrange^+^ (**Figure 4A’’’-A’’’’**), confirming that the reporter line allows the visualization of *NvPrdm14d*^+^ NPCs and their progeny.

**Figure 4:**
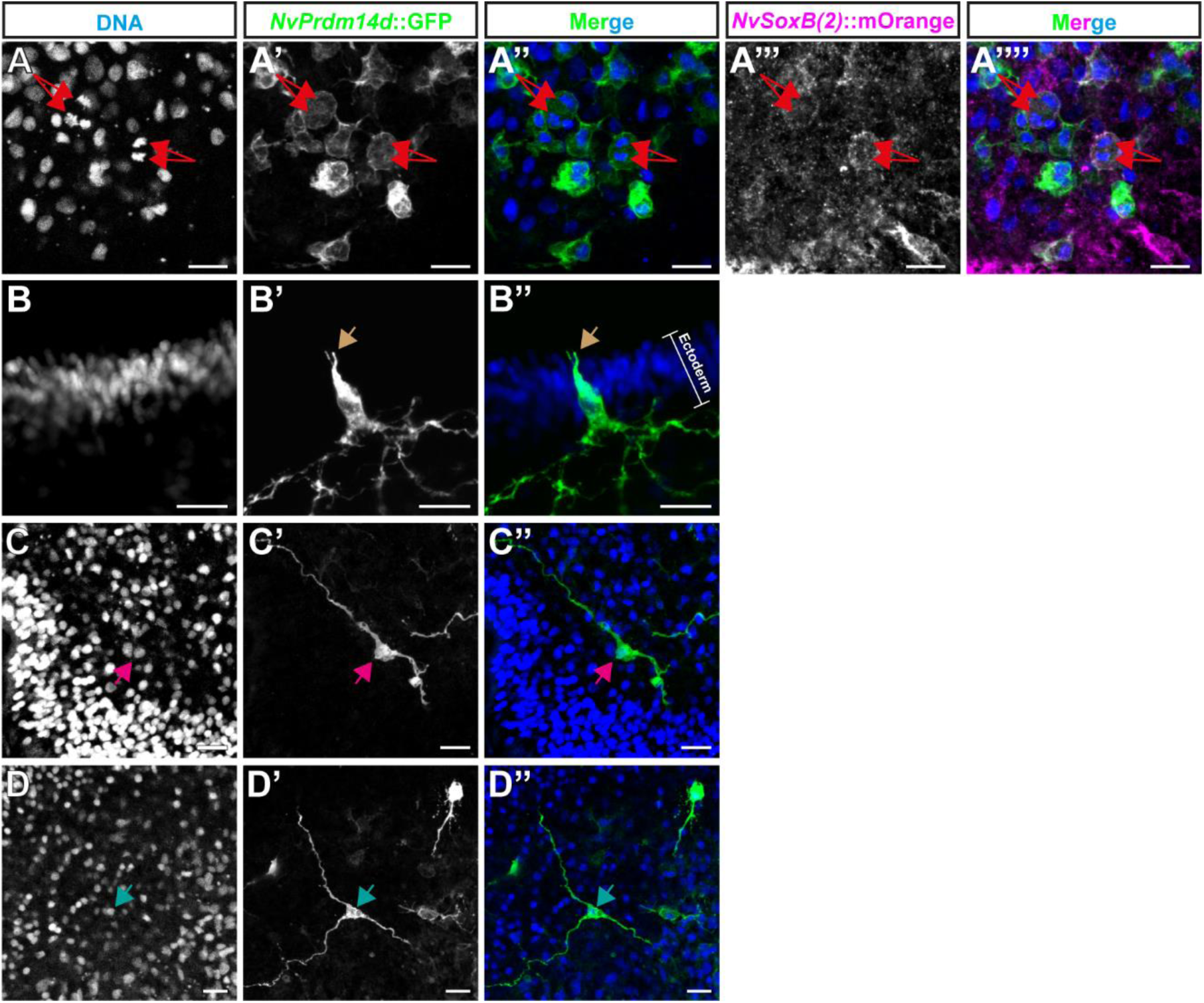
Cells expressing NvPrdm14d::GFP are mainly endodermal ganglion neurons. Overview of the different types of cells labelled by NvPrdm14d::GFP, shown in green. (**A**) NvPrdm14d::GFP labels dividing endodermal neural progenitor cells (NPCs) as these cells are co-labelled by NvSoxB(2)::mOrange, shown in magenta. Arrows indicate sister chromatids at anaphase. (**B**) Consistent with the expression pattern of NvPrdm14d, some rare ectodermal sensory neurons are labelled by NvPrdm14d::GFP in the central region of the oral-aboral axis. Arrows indicate the sensory cilium; the white bracket indicates the ectoderm. (**C-D**) The endodermal neurons labeled by NvPrdm14d::GFP are ganglion cells with a variable number of neurites, e.g. bi- and tripolar neurons as depicted here. No asymmetric distribution of these ganglion neurons was observed along the embryo axes. Arrows indicate neural soma. Scale bars: 10 µm

In late planulae, we could observe some rare ectodermal neurons labelled by GFP. Only few of the screened larvae exhibited such cells (12%, n=42) and only few ectodermal cells were found in individual larvae (between 1 and 3 cells). These cells are likely derived from the few ectodermal cells expressing *NvPrdm14d* as revealed by ISH (**Figure 1**). These GFP^+^ ectodermal neurons are found in the central region of the oral-aboral axis, are elongated along the apical-basal axis of the ectoderm and display an apical cilium (**Figure 4B-B’’**). The morphological characteristics of these cells suggests that they are sensory neurons.

By contrast, most GFP^+^ neurons in planula larvae have their soma located at a basal position within the endodermal epithelium, suggesting that they are ganglion neurons (**Figure 4C-D’’**). These endodermal neurons exhibit a variable number of neurites, hence can be identified as bipolar and tripolar neurons. However, no specific spatial distribution of these neurons was detected.

These results show that the *NvPrdm14d* reporter labels a subset of the endodermal NPCs as well as their progeny, which mainly consists of ganglion neurons.

### *NvPrdm14d*^+^ cells generate a small subset of endodermal neurons

To further investigate the identity of the endodermal neurons generated by *NvPrdm14d*^+^ NPCs, we crossed the transgenic line with other reporter lines that have been characterized previously.

The *NvElav1::mOrange* reporter line has been shown to label a large population of differentiated neurons in both the ectoderm and the endoderm, but not cnidocytes (Nakanishi et al., 2012). Though many *NvPrdm14d*::GFP^+^ neurons and neurites are in close proximity of *NvElav1*::mOrange^+^ neurons in primary polyps, we did not detect any co-expression of GFP and mOrange in both planula larvae and primary polyps (**Figure 5A-F’’**). This was unexpected since *NvPrdm14d* and *NvElav1* are partially co-expressed in the DFISH experiment (**Figure 2E-H**). Neither of the transgenic reporter lines matches the expression of the endogenous gene perfectly (Nakanishi et al., 2012) (**Figure 3**) and we assume that this is reflected in the lack of co-expression in the double transgenics. Together, these data show that the *NvElav1*::mOrange and *NvPrdm14d*::GFP lines label two distinct populations of neurons.

**Figure 5:**
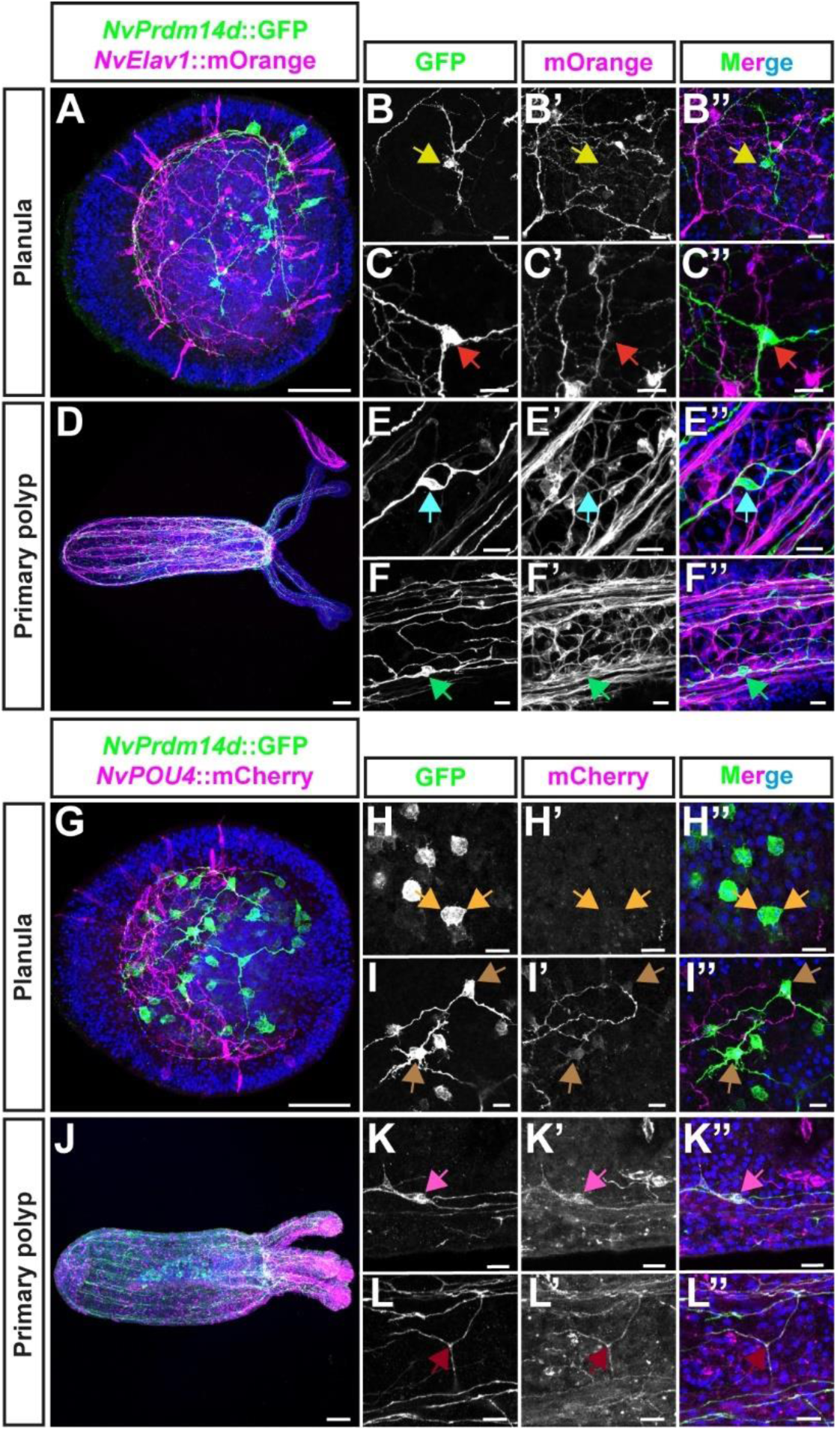
NvPrdm14d::GFP highlights a subset of endodermal NvPOU4::mCherry^+^ neurons but not of NvElav1::mOrange+ neurons. (**A-F**) Confocal images of immunofluorescence staining for NvPrdm14d::GFP and NvElav1::mOrange, respectively shown in green and magenta. (**A-C**) Planula stage. (**D-F**) Primary polyp stage. (**B-C & E-F**) Enlargements showing that endodermal neurons expressing NvPrdm14d::GFP do not express for NvElav1::mOrange. Arrows indicate the soma of NvPrdm14d::GFP^+^ neurons that do not express NvElav1::mOrange. (**G-L**) Confocal images of immunofluorescence staining for NvPrdm14d::GFP and NvPOU4::mCherry, respectively shown in green and magenta. (**G-I**) Planula stage. (**J-L**) Primary polyp stage. (**H**) Enlargements showing dividing NPCs expressing NvPrdm14d::GFP but not NvPOU4::mCherry, consistent with the role of NvPOU4 in terminal differentiation. Arrows indicate chromosomes at anaphase in NvPrdm14d::GFP^+^ NPCs that do not express NvPOU4::mCherry. (**I & K-L**) Enlargements showing that differentiated neurons expressing NvPrdm14d::GFP also express NvPOU4::mCherry. Arrows indicates neurons or neurites co-expressing NvPrdm14d::GFP and NvPOU4::mCherry. Embryos are counterstained for DNA in blue, the oral pole is oriented to the right, scale bars: 50 µm in (**A, D, G & J**) and 10 µm in (**B-C, E-F, H-I & K-L**).

Next, we crossed the *NvPrdm14d::GFP* line with the *NvPOU4::mCherry* reporter line that has been shown to label ectodermal and endodermal neurons, as well as cnidocytes in *Nematostella* (Tournière et al., 2020). At planula stage, we observed a partial co-expression of GFP and mCherry (**Figure 5G-I’’**), which matches our observation that *NvPrdm14d* and *NvPOU4* transcripts are partially co-expressed (**Figure 2I-L**). Among the cells co-labelled by both transgenes, we did not observe any of the *NvPrdm14d*::GFP^+^ cells that lack neurites or divide (**Figure 5H-H’’**). However, *NvPrdm14d*::GFP^+^ cells exhibiting neurites always co-express *NvPOU4*::mCherry (**Figure 5I-I’’**), although frequently at comparably low levels. This is in line with the role of *NvPOU4* in terminal differentiation of neurons (Tournière et al., 2020). Furthermore, we noted that the *NvPrdm14d*::GFP^+^ endodermal neurons co-express the *NvPOU4*::mCherry reporter in the primary polyp, but that the majority of *NvPOU4*::mCherry^+^ neurons are *NvPrdm14d*::GFP^−^ (**Figure 5J-L’’**).

The analysis of these double transgenic animals reveals that the *NvPrdm14d*::GFP and *NvPOU4*::mCherry reporters are co-expressed in a population of post-mitotic neurons and that the *NvPrdm14d*^+^ NPCs likely generate only a small subset of endodermal neurons.

### The *NvPrdm14d*::*GFP* transgene highlights a population of neurons in the close vicinity of retractor muscles

A role for *Prdm14* in neurogenesis has been described in zebrafish where it is involved in the axon outgrowth of primary motoneurons (Liu et al., 2012). We therefore decided to explore the possibility that *NvPrdm14d* might be involved in the development of motoneurons in *Nematostella*.

To this end, we crossed the *NvPrdm14d::GFP* reporter line with the *NvMyHC1::mCherry* reporter line, specifically labelling retractor and longitudinal tentacle muscles through the expression of mCherry under the *Myosin Heavy Chain-ST* promoter (Renfer et al., 2010). In primary polyps, we observed that some of the *NvPrdm14d*::GFP^+^ neurons are near the retractor muscles (**Figure S2A-C**), which are located on endodermal infoldings of the body wall called mesenteries (**Figure 6A-A’’**). However, the developing retractor muscles are very close to the parietal muscle and the longitudinal neurite tracts of the body wall at this stage, confounding the analysis of a potential association with *NvPrdm14d*::GFP^+^ neurons.

**Figure 6:**
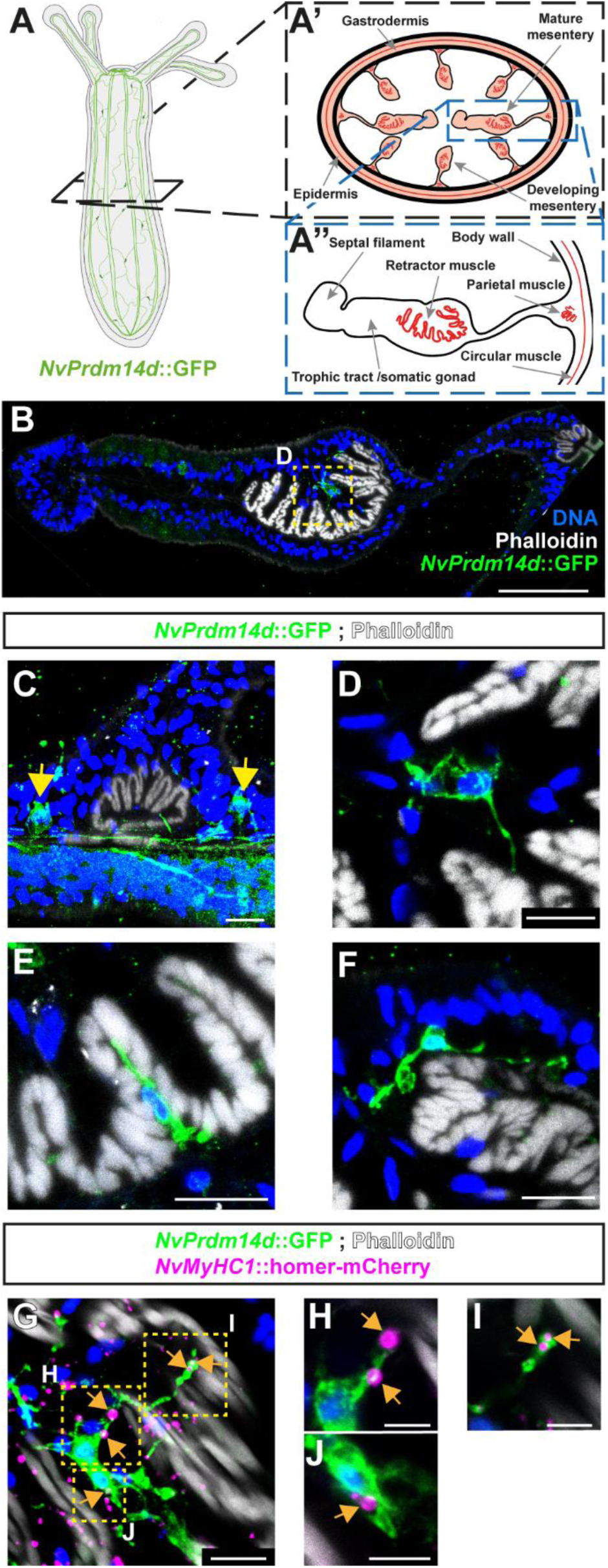
NvPrdm14d::GFP^+^ neurons are in the close vicinity of retractor muscles. (**A**) Schematic illustrating the experimental sectioning performed to observe NvPrdm14d::GFP^+^ neurons in the vicinity of retractor muscles. The different structures observed on sections and in mesenteries are described in (**A’-A’’**). (**B-J**) Confocal images of immunofluorescence staining for NvPrdm14d::GFP in mesenteries. GFP is shown in green and muscles in white. *(C)* Enlargement of a parietal muscle showing a concentration of neurites on both sides (arrows), corresponding to the longitudinal tracts. (**D-F**) Enlargements of different neurons expressing the NvPrdm14d::GFP in the vicinity of the retractor muscles. *(D)* Enlargement corresponding to the dashed line square shown in (**B**). (**G-J**) NvPrdm14d::GFP expressing neurons project neurites toward retractor muscles and are in contact with putative neuromuscular synaptic junctions. The putative neuromuscular synaptic junctions are revealed by a fusion protein homer- mCherry under the control of the retractor muscle- specific promoter NvMyHC1, shown in magenta. Arrows indicate putative neuromuscular post- synapses in contact with NvPrdm14d::GFP^+^ neurons/neurites. (**H-J**) Single images extracted from the projection shown on G. They depict enlargements of close proximity between NvPrdm14d::GFP^+^ cells and putative neuromuscular synaptic junctions. Samples are counterstained for DNA in blue. Scale bars: 50 µm in (**B**), 10 µm in (**C-G**) and 5 µm in (**H-J**).

Therefore, we decided to image 2 to 4 months-old polyps with well-developed mesenteries (**Figure 6**). The analysis of mesentery cross-sections stained with phalloidin (to visualize F-actin) revealed a concentration of *NvPrdm14d*::GFP^+^ neurites, likely corresponding to the longitudinal neurite tracts, as well as some cell bodies located on each side of the parietal muscles (**Figure 6C**). We also observed some *NvPrdm14d*::GFP^+^ neurons within the mesenteries in close vicinity of retractor muscles (**Figure 6B & D-F**). We did not find *NvPrdm14d*::GFP^+^ neurons in any other part of the mesenteries. Interestingly, these neurons were observed at diverse locations around the retractor muscles. Some are close to or within the endodermal epithelium (**Figure 6D, F**), others are located on the basal side of the retractor muscles, likely within the mesoglea (**Figure 6E**). All these neurons appear to project neurites either towards the contractile filaments of the retractor muscles or along them, suggesting that they might establish synaptic connections with these muscles.

To test this hypothesis, we generated a *NvMyHC1::homer-mCherry* line expressing the putative post-synaptic protein homer, fused to mCherry, under the control of the retractor muscle- specific *NvMyHC1* promoter, presumably allowing the visualization of post-synaptic sites in the retractor muscles. Indeed, the homer-mCherry fusion protein is detected in puncta arranged in longitudinal tracts in the body column and tentacles of primary polyps (**Figure S3A-E**), consistent with the expression of the mCherry protein labelling retractor and tentacle muscles in *NvMyHC1::mCherry* primary polyps [**Figure S3L-P**, (Renfer et al., 2010)]. Following cross- sectioning of *NvMyHC1*::homer-mCherry^+^ polyps, we observed that puncta are found close to the contractile filaments of the retractor muscles as revealed by phalloidin staining, both in mature and in developing mesenteries (**Figure S3H-K**). We did not observe such puncta in *NvMyHC1*::mCherry polyps (**Figure S3H-K & S-V**).

We then analyzed cross-sections of *NvPrdm14d::GFP*; *NvMyHC1::homer-mCherry* double transgenic animals. We observed that somata and neurites of the *NvPrdm14d*::GFP^+^ neurons located in the vicinity of retractor muscles are in close contact with *NvMyHC1*::homer-mCherry^+^ puncta (**Figure 6G-J**). This observation supports the hypothesis that *NvPrdm14d*::GFP^+^ neurons establish synaptic connections with retractor muscles.

We did not observe any *NvPrdm14d*::GFP^+^ neurons or neurites in the tissue connecting the body wall and the retractor muscles (**Figure 6B**). When we analyzed cross-sections of *NvPrdm14d::GFP*; *NvElav1::mOrange* double transgenic polyps, we could not detect *NvElav1*::mOrange^+^ neurons between the body wall and the retractor muscle (**Figure S2D**). The existence of an additional population of neurons, not labelled by the two transgenes, is a possible explanation for this observation.

Together, these data reveal that the *NvPrdm14d*::GFP reporter highlights a population of previously undescribed potential motoneurons in the vicinity of retractor muscles.

### *NvPrdm14d*^+^ cells do not originate in the ectodermal pharynx

Since the expression pattern of *NvPrdm14d* is dynamic during embryonic development with a strong expression in the pharynx before being mainly being expressed in scattered endodermal cells (**Figure 1**), we wanted to take advantage of the *NvPrdm14d::GFP* transgenic line to confirm the endodermal origin of the *NvPrdm14d*^+^ cells. Indeed, the part of the pharynx lining the oral opening has an ectodermal origin, hence it is possible that the *NvPrdm14d*^+^ NPCs originate in this domain before migrating to the endoderm at planula stage.

To address this possibility, we used a transgenic reporter line expressing mOrange2 under the control of the ectodermal, pharynx-specific 5.9kb long *NvFoxA* promoter (Fritzenwanker et al., 2004; Martindale et al., 2004; Steinmetz et al., 2017), and crossed it with the *NvPrdm14d::GFP* line. At early planula stage, the *NvFoxA*::mOrange2 reporter is exclusively expressed in the ectodermal part of the pharynx (**Figure 7A**). In such larvae, we observed that few *NvPrdm14d*::GFP^+^ cells are neighboring *NvFoxA*::mOrange2^+^ cells, but none of them express both reporters (**Figure 7B-B’’**). By contrast, most of *NvPrdm14d*::GFP^+^ cells in the pharynx are distant from *NvFoxA*::mOrange2^+^ cells. Moreover, none of the *NvPrdm14d*::GFP^+^ differentiated neurons express *NvFoxA*::mOrange2 (**Figure 7C-C’’**). Similar observations were made in late planula larvae that exhibit a broader expression of *NvFoxA*::mOrange2 in the ectodermal pharynx with a higher number of *NvFoxA*::mOrange2^+^ cells (**Figure 7D-F’’**). In the primary polyp, *NvFoxA*::mOrange2^+^ cells are found in the mesenteries where they form the distal structures called septal filaments (**Figure 7G**), consistent with a previous report (Steinmetz et al., 2017). Similar to our description in planula larvae, we did not detect any *NvFoxA*::mOrange2^+^ cells in the ectodermal pharynx or the septal filaments, co-expressing *NvPrdm14d*::GFP (**Figure 7H-I’’**). At this stage, none of the *NvPrdm14d*::GFP^+^ differentiated neurons co-express *NvFoxA*::mOrange2 either (**Figure 7J-J’’**).

**Figure 7:**
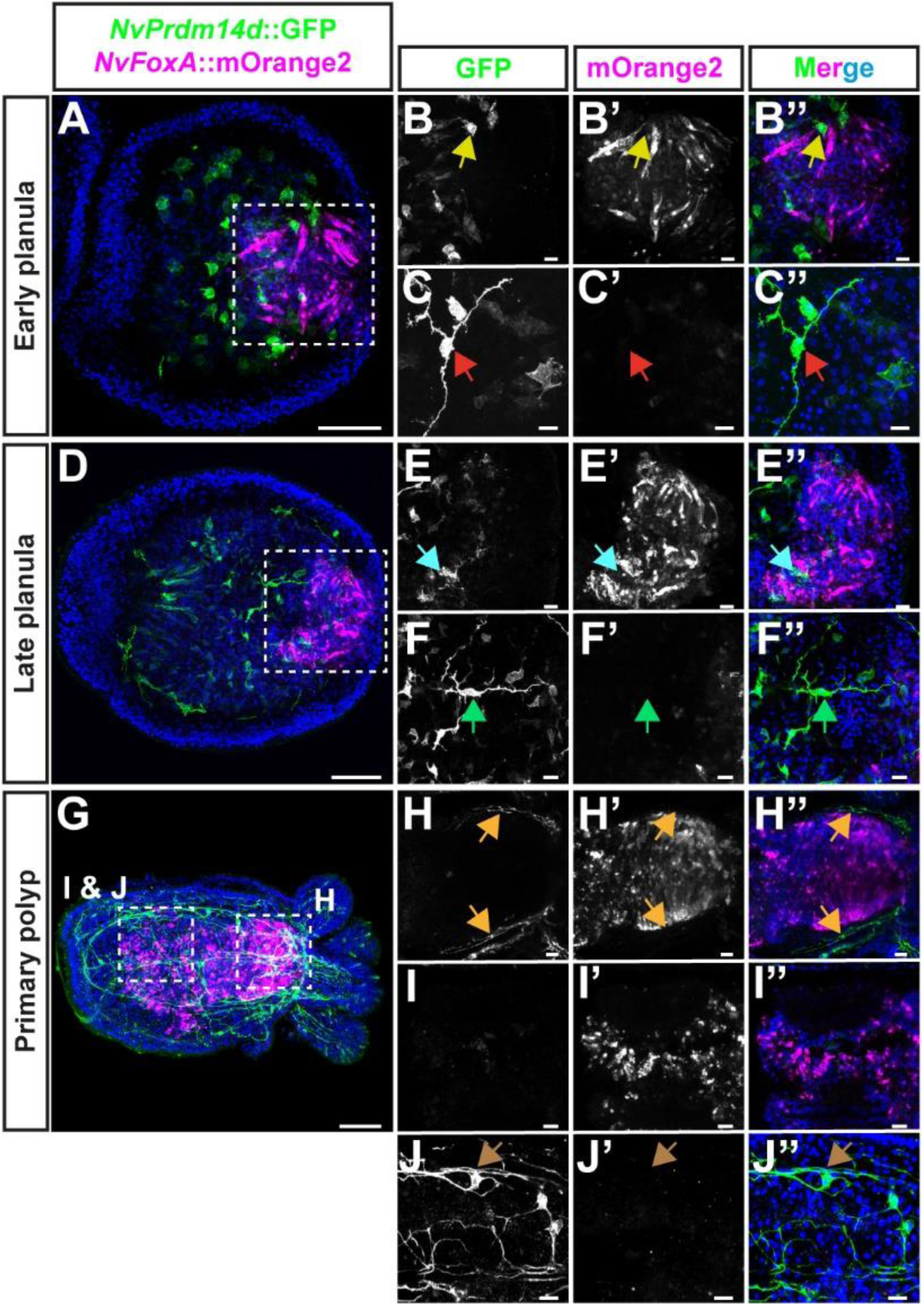
NvPrdm14d::GFP^+^ neurons do not originate in the ectodermal pharynx. (**A-J**) Confocal images of immunofluorescence staining for NvPrdm14d::GFP and NvFoxA::mOrange2, respectively shown in green and magenta. The promoter of NvFoxA drives gene expression in the ectodermal pharynx, specifically. (**A-F**) Planula stages. (**G-J**) Primary polyp stage. (**B, E & H**) Enlargements showing that ectodermal pharyngeal cells express NvFoxA::mOrange2 but not NvPrdm14d::GFP. Arrows indicate that NvPrdm14d::GFP^+^ neurons are negative for NvFoxA::mOrange2. (**I**) Enlargements showing ectodermal pharyngeal cells in the distal part of the mesenteries of primary polyps. Those cells express NvFoxA::mOrange2 but not NvPrdm14d::GFP. (**C, F & J**) Enlargements showing that endodermal neurons expressing NvPrdm14d::GFP, do not express NvFoxA::mOrange2. Arrows indicate the soma of some NvPrdm14d::GFP^+^ neurons showing an absence of NvFoxA::mOrange2 expression. Embryos are counterstained for DNA in blue, the oral pole is oriented to the right, scale bars: 50 µm in (**A, D & G**) and 10 µm in (**B-C, E-F & H-J**).

Taken together, these observations support the conclusion that endodermal *NvPrdm14d*^+^ cells originate from the endoderm, including the endodermal region of the pharynx, and not from ectodermal pharyngeal cells.

### The transcriptome of *NvPrdm14d*::GFP^+^ cells provides new candidate genes involved in endodermal neurogenesis

To gain further insights into the characteristics of neurons derived from *NvPrdm14d*^+^ NPCs, we separated *NvPrdm14d*::GFP^+^ and *NvPrdm14d*::GFP^-^ (control) cells using fluorescence-activated cell sorting (FACS) at primary polyp stage and performed RNA sequencing on both cell populations in triplicates (**Figure 8A-B**).

**Figure 8:**
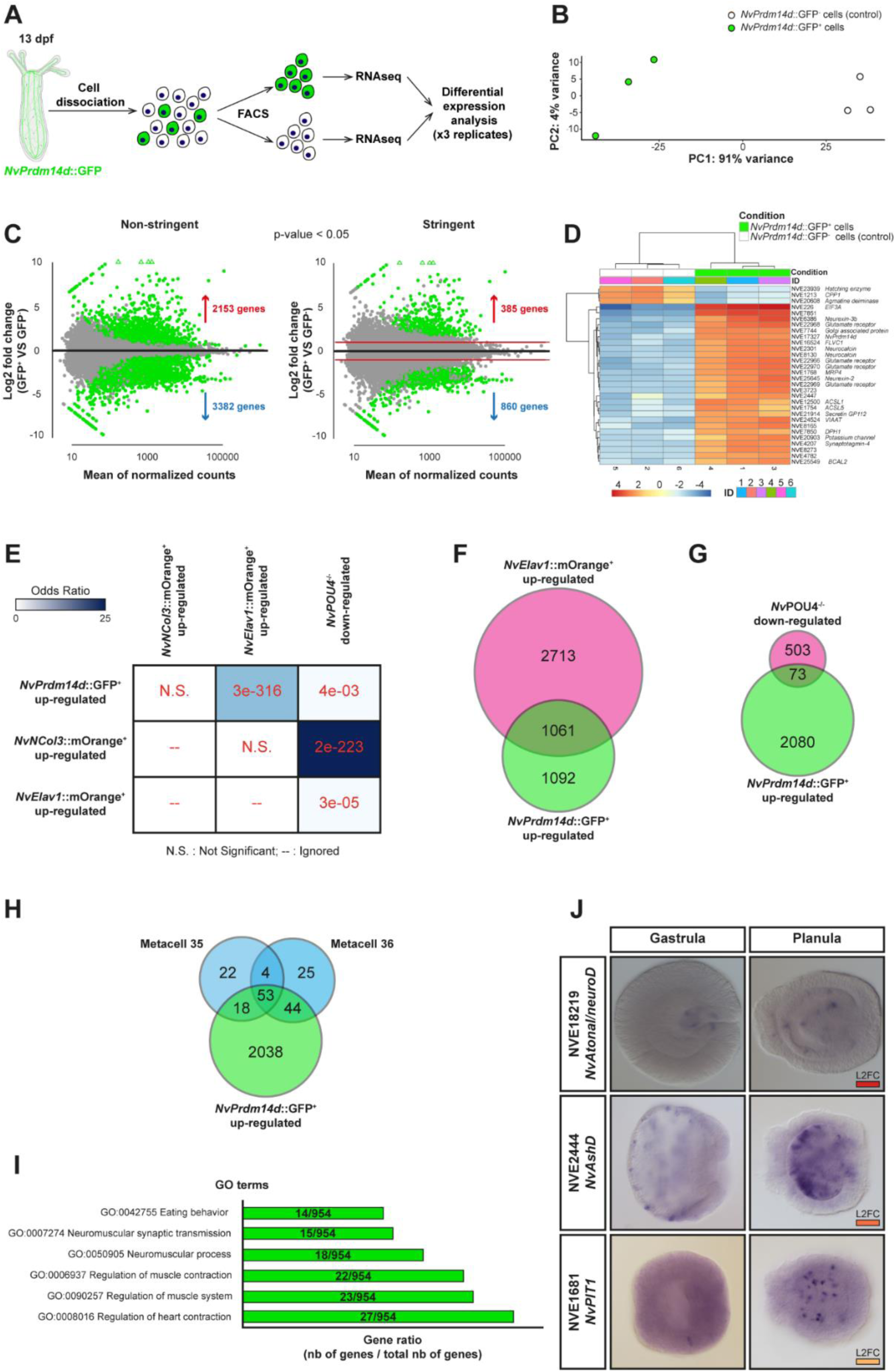
Transcriptomic analysis of NvPrdm14d::GFP^+^ cells. (**A**) Schematic illustrating the experimental design. NvPrdm14d::GFP^+^ cells are green and NvPrdm14d::GFP^-^ cells are white. (**B**) PCA plot using the gene counts after variance stabilizing transformation (VST). Green dots represent NvPrdm14d::GFP^+^ samples and white dots represent control NvPrdm14d::GFP^-^ samples. (**C**) MA-plots of up- and downregulated genes between NvPrdm14d::GFP^+^ cells and NvPrdm14d::GFP^-^ controls. Differentially expressed genes are shown in green with a p-value<0.05. The plot on the right shows differentially expressed genes with a stringent filter using a more conservative test and excluding genes with a log2 fold change comprised between -1 and 1. (**D**) Gene clustering matrix showing a subset of the most highly variable genes explaining the difference between NvPrdm14d::GFP^+^ and NvPrdm14d::GFP^-^ cells. Notice that NvPrdm14d appears there. (**E**) Comparison of the overlap between non-stringent upregulated genes in NvPrdm14d::GFP^+^ cells with those upregulated in NvElav1::mOrange^+^ and NvNCol3::mOrange^+^ cells; as well as with those downregulated in NvPOU4 mutants. The comparison is done using the GeneOverlap R package. The strength of the blue color indicates the odds ratio and numbers indicate the p-value, calculated using Fisher’s exact test. (**F**) Venn diagram comparing the number of non-stringent genes upregulated in NvPrdm14d::GFP^+^ cells with those upregulated in NvElav1::mOrange^+^ cells. (**G**) Venn diagram comparing the number of non- stringent genes upregulated in NvPrdm14d::GFP^+^ cells with those down-regulated in NvPOU4 mutants. (**H**) Venn diagram comparing the number of non-stringent genes upregulated in NvPrdm14d::GFP^+^ cells with those characterizing the neuronal metacells 35 and 36 from the single cell transcriptome of whole Nematostella by (Sebé-Pedrós et al., 2018). (**I**) GO terms, associated with motoneuron function, found among non-stringent genes upregulated in NvPrdm14d::GFP^+^ cells. The gene ratio shows the number of genes associated with these terms in NvPrdm14d::GFP^+^ cells, compared to the total number of genes associated with GO terms in NvPrdm14d::GFP^+^ cells. (**J**) Colorimetric in situ hybridization of a sample of stringent genes upregulated in the NvPrdm14d::GFP^+^ cells showing an expression pattern consistent with a role in the development of endodermal neurons. Note that NvAtonal/neuroD is not upregulated in NvElav1::mOrange^+^ cells. L2FC: Log2 fold change, the strength of the red color indicates the level of up-regulation as shown in (**D**).

In the *NvPrdm14d*::GFP^+^ cells, a total of 5,535 genes are differentially expressed, including 2,153 enriched and 3,382 depleted genes (“non-stringent list”, p<0.05, **Figure 8C left**). Additionally, we generated a “stringent list” by removing all differentially expressed genes with a Log2Fold Change between -1 and 1. This allowed the identification of 1,245 differentially expressed genes, including 285 enriched and 860 depleted genes (p<0.05, **Figure 8C right**). Within these lists, we found *NvPrdm14d* as one of the most differentially expressed genes supporting the notion that *NvPrdm14d* remains expressed in at least some of the GFP^+^ neurons at least until primary polyp stage (**Figure 8D**).

First, we looked at the non-stringent list and sought for genes that would help characterizing the neural features of *NvPrdm14d*::GFP^+^ cells. We found many genes annotated as receptors for glutamate, neuropeptide FF, pyroglutamylated RFamide, acetylcholine and dopamine (**Table S2**). We also found some genes annotated as receptors for adrenaline and glycine. Additionally, we observed an enrichment in calcium-, sodium- and potassium-gated channels. Moreover, we identified several genes encoding putative pre-synaptic proteins, such as synaptotagmins (**Table S2**).

Then, we analyzed whether target genes and/or genes encoding functional or physical interactors of Prdm14 known from other species, were found among the upregulated genes in *NvPrdm14d*::GFP^+^ cells. We found that *NvIslet* (NVE10444), a target of Prdm14 in zebrafish motoneurons, is upregulated in *NvPrdm14*::GFP^+^ cells (stringent list, **Table S2**). In addition, we found the Prdm14 partners *NvCBFA2T* (NVE5561) and *NvEed-A* (NVE21521, part of PRC2) in the non-stringent list, indicating that the Prdm14-PRC2 repressor complex might be conserved in *Nematostella* and involved in the development of *NvPrdm14d*::GFP^+^ neuron (**Table S2**).

Next, we compared the *NvPrdm14d*::GFP^+^ transcriptome with previously generated transcriptomes, *i.e.* for *NvNCol3*::mOrange2^+^ and *NvElav1*::mOrange^+^ cells (Gahan et al., 2022; Tournière et al., 2020). All three transcriptomes were generated from primary polyps at 13 dpf (days post-fertilization). For each comparison, we used the GeneOverlap R package (Shen and Sinai, 2020) to test whether the gene overlap is random or not. This test provides the odds ratio (strength of association) and its p-value (significance of the association). An odds ratio equal or below 1 means no association, while the association is stronger with a higher ratio. We found a small overlap between genes upregulated in *NvPrdm14d*::GFP^+^ cells (∼14.3%) and those upregulated in *NvNCol3*::mOrange2^+^ cells, however the odds ratio indicates that this overlap is not significant (**Figure 8E**).

By contrast, we found that about 50% of the genes upregulated in *NvPrdm14d*::GFP^+^ cells are also upregulated in *NvElav1*::mOrange^+^ cells, and this overlap is significant (**Figure 8E-F**). We also found *NvElav1* among the upregulated genes (stringent list, **Table S2**). This is in line with the co-expression of *NvPrdm14d* and *NvElav1* as shown by our DFISH experiment (**Figure 2E-H**) and it suggests that the *NvElav1::mOrange* transgenic reporter line does not label the entirety of *NvElav1*^+^ cells.

As we observed co-expression of *NvPrdm14d* and *NvPOU4* (**Figure 2I-L**), and of the *NvPrdm14d*::GFP and *NvPOU4*::mCherry reporters (**Figure 5G-I’’**), we compared our transcriptome with the one of *NvPOU4* mutants, referred to as *NvPOU4*^-/-^ (Tournière et al., 2020). We found a small, but significant, overlap between genes upregulated in *NvPrdm14d*::GFP^+^ cells (∼3.4%) and those downregulated in *NvPOU4*^-/-^ (**Figure 8E & G**).

Therefore, *NvPOU4* might regulate aspects of the terminal differentiation of *NvPrdm14d*^+^ neurons.

Furthermore, we took advantage of the whole-animal single cell transcriptome atlas of *Nematostella* and searched for neuronal metacells expressing *NvPrdm14d* (Sebé-Pedrós et al., 2018). We found two metacells (35 & 36) expressing *NvPrdm14d* and they respectively share 73.5% and 77.2% of their defining genes with genes upregulated in the *NvPrdm14d*::GFP^+^ transcriptome (**Figure 8H, Table S3**). Moreover, these two metacells share about 50% of their defining genes (**Figure 8H**). However, we did not find such an overlap in gene expression when we analyzed genes upregulated in the *NvPrdm14d*::GFP^+^ transcriptome within all the neuronal metacells, confirming the specific expression of *NvPrdm14d* in the metacells 35 and 36.

Altogether, these data support the hypothesis that *NvPrdm14d* is expressed in, at least, two neural cell types.

With the aim to understand unique characteristics of *NvPrdm14d*::GFP^+^ neurons, we next looked at genes that are upregulated in these neurons, but not upregulated in *NvElav1*::mOrange^+^ cells. This revealed an enrichment in GO terms associated with chromosome and chromatin organization, as well as cell cycle regulation (**Table S4**). While this is not informative about the potential role played by *NvPrdm14d*::GFP^+^ neurons, it is consistent with the fact that *NvPrdm14d*-expressing cells include NPCs, whereas *NvElav1* is exclusively expressed in post-mitotic neurons. We then specifically sought for GO terms associated with motoneuron functions as *NvPrdm14d*::GFP^+^ neurons potentially establish synaptic connections with retractor muscles. We found six GO terms related to neuromuscular functions, however they are not the most enriched terms (**Figure 8I**). Thus, the GO term analysis does not provide unambiguous support for a potential motoneuron identity among the *NvPrdm14d*::GFP^+^ neurons.

Finally, we focused on transcription factors upregulated in *NvPrdm14d*::GFP^+^ cells and we checked their expression pattern in the literature, when existing. We found that most of the genes whose expression pattern is available, are expressed in the endoderm. These genes are *NvDmrt-b* [NVE6455, (Bellefroid et al., 2013; Parlier et al., 2013)], *NvGATA* [NVE8199, (Martindale et al., 2004; Steinmetz et al., 2017)], *NvGCM* [NVE12024, (Marlow et al., 2009; Marlow et al., 2012)], *NvIslet* [NVE10444, (Steinmetz et al., 2017)], *NvNkx2*.2D [NVE10557, (Steinmetz et al., 2017)], *NvPOU4* [NVE5471, (Tournière et al., 2020)], *NvNkx3* [NVE18255, (Marlow et al., 2013; Steinmetz et al., 2017)], *NvTbx20.3* [NVE24169, (Steinmetz et al., 2017)] and *NvSix1/2a* [NVE9850, (Steinmetz et al., 2017)]. Although some of these genes are additionally expressed in the ectoderm, we noticed that expression of *NvGCM*, *NvNkx2.2D*, *NvTbx20.1* and *NvSix1/2a* appears to be restricted to the endoderm, suggesting a role for these genes specifically in endodermal neurogenesis. ISH for some of the previously uncharacterized transcription factors enriched in the *NvPrdm14d*::GFP^+^ cells showed that *NvAtonal/neuroD* (NVE18219), *NvAshD* (NVE2444) and *NvPIT1* (NVE1681) are mainly expressed in scattered endodermal cells at planula stage, although they display expression in some ectodermal cells at gastrula stage (**Figure 8J**). Interestingly, *NvAtonal/neuroD* was not found in genes upregulated in *NvElav1*::mOrange^+^ cells, indicating that this gene might be exclusively expressed in the same lineage as *NvPrdm14d*. This gene is expressed in the pharynx of gastrula larvae, and then in scattered endodermal cells in planula larvae, though in fewer cells than *NvPrdm14d* (**Figure S4**), suggesting that it might represent a subpopulation of *NvPrdm14d*::GFP^+^ neurons.

Taken together, the *NvPrdm14d*::GFP^+^ transcriptome analysis showed that *NvPrdm14d*::GFP^+^ neurons likely are a subpopulation of the *NvPOU4* endodermal nervous system, that they overlap at least partially with *NvElav1*^+^ neurons and that they include putative motoneuron-like cells.

### *NvPrdm14d* is a positive and negative regulator of gene expression

To determine whether genes upregulated in *NvPrdm14d*::GFP^+^ cells are regulated by *NvPrdm14d*, we performed knockdown experiments by injection of shRNAs (He et al., 2018). We designed two shRNAs that reduced *NvPrdm14d* transcript levels to 18.6% (log2 fold change -2.43) and 30.5% (log2 fold change -1.71), respectively, compared to animals injected with a control shRNA targeting GFP (**Figure 9**). We selected four genes upregulated in *NvPrdm14d*::GFP^+^, but not in *NvElav1*::mOrange^+^ cells and analyzed their expression by quantitative RT-PCR. We found that three genes putatively related to neural differentiation and function [*NVE25645* (neurexin-related), *NVE24524* (putative vesicular inhibitory amino acid transporter) and *NVE22970* (putative ionotropic glutamate receptor)] were downregulated in planula injected with *NvPrdm14d* shRNAs (**Figure 9**). In contrast, the expression of *NvAtonal/neuroD* was upregulated (log2 fold change 1.30 and 0.59, respectively, for the two shRNAs, **Figure 9**).

**Figure 9:**
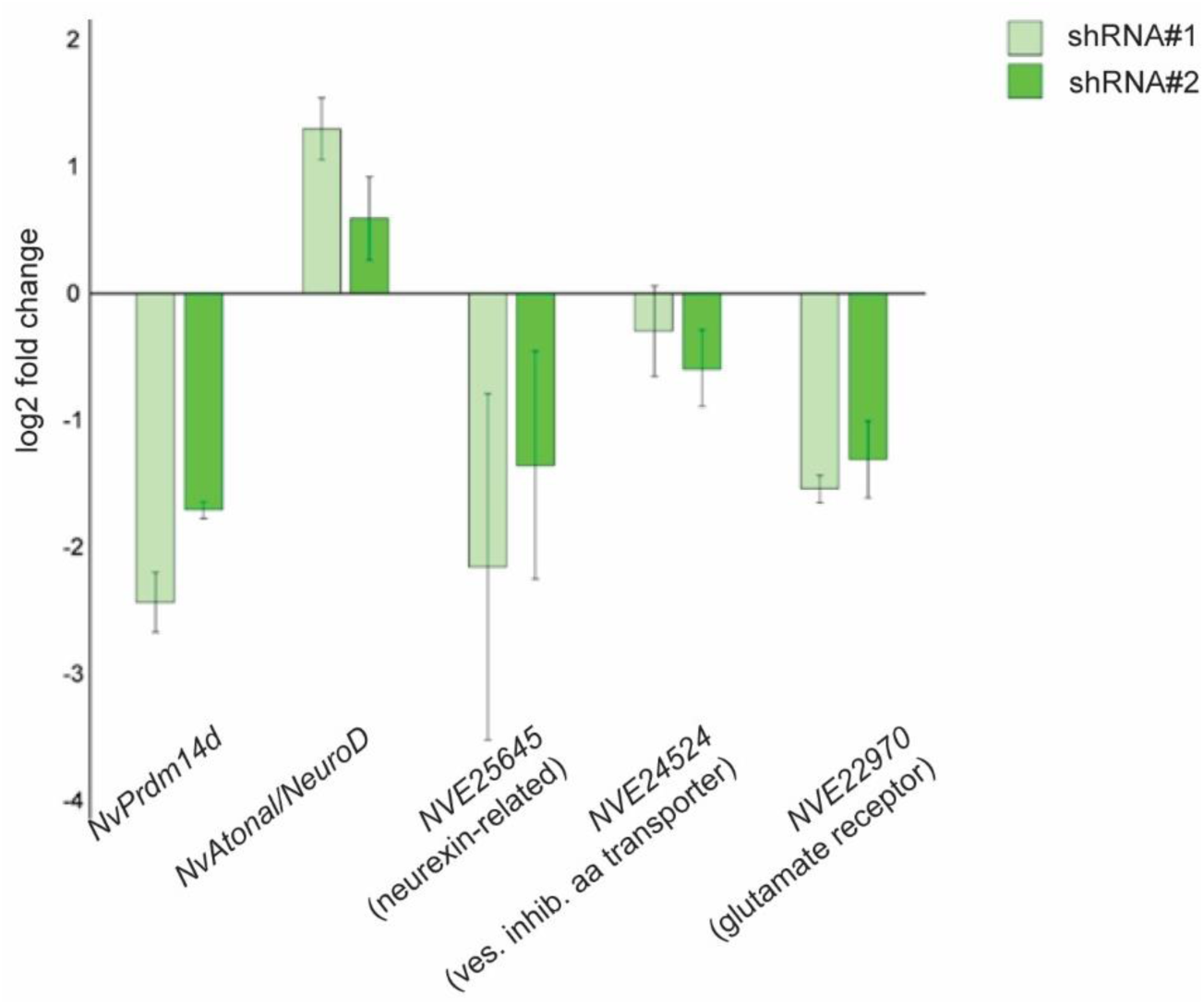
NvPrdm14d is a positive and negative regulator of gene expression. qRT-PCR on selected neural genes upregulated in NvPrdm14d::GFP^+^, but not in NvElav1::mOrange^+^ cells, at early planula stage after shRNA injection. The graph shows the mean ± SD of the log2 fold change in expression between control shRNA and either NvPrdm14d shRNA #1 (light green) or #2 (green) across three independent biological replicates.

These results show that *NvPrdm14d* has both positive and negative effects on gene expression and is likely required for the acquisition of at least some neural features.

## DISCUSSION

In this report, we find that all four *NvPrdm14* paralogs are expressed in the nervous system of *Nematostella vectensis*. We show that the expression of *NvPrdm14d* defines a subpopulation of endodermal NPCs generating a subset of the endodermal nervous system, including potential motoneurons in *Nematostella*. The overlap with *NvPOU4::mCherry* expression suggests that their terminal differentiation might be, in part, regulated by *NvPOU4*. Through the analysis of the *NvPrdm14d*::GFP^+^ transcriptome, we identified a panel of genes potentially involved in endodermal neurogenesis in *Nematostella*, either specifically in the *NvPrdm14d* lineage or more broadly. Finally, our functional analysis of *NvPrdm14d* revealed that this gene has positive and negative effects on neural gene expression in *Nematostella*.

### Endodermal origin of *NvPrdm14d*^+^ neurons and the heterogeneity of NPCs

The expression of *NvPrdm14d* is initially detectable in the pharynx before expression in scattered cells throughout the endoderm becomes visible (**Figure 1**). One explanation for these dynamics would be a migration of neural progenitors or precursors from the pharynx into the body wall endoderm. Since the pharynx consists of an ectodermal and endodermal part, the *NvPrdm14d*^+^ neurons in the body wall endoderm could potentially be of ectodermal origin. Two observations argue against this scenario. First, *NvPrdm14d* expression is predominantly found in the endodermal part of the pharynx (**Figure 2M**) and second, the *NvPrdm14d*::GFP^+^ cells in the body wall endoderm do not express *NvFoxA*::mOrange2 (**Figure 7**), which labels the ectodermal pharynx and cells derived from it. This suggests that the *NvPrdm14d*^+^ neurons are indeed of endodermal origin.

Previous work has shown that ectodermal and endodermal neurogenesis in *Nematostella* are both regulated by transcription factors [*NvAth-like* and *NvSoxB(2)*] and signaling molecules (Notch) that act at the level of neural/secretory progenitor cells [N/SPCs, (Layden and Martindale, 2014; Richards and Rentzsch, 2014; Richards and Rentzsch, 2015)]. The expression of *NvPrdm14d* in a subpopulation of *NvSoxB(2)*^+^ endodermal N/SPCs (**Figure 2**) that gives rise only to neurons, shows that progenitor cell diversity, and not only terminal differentiation, is part of the endoderm-specific neurogenic program in *Nematostella.* We speculate that other populations of endodermal neurons derive from molecularly distinct progenitor cells and that the putative different types of neural progenitors can be considered as intermediate progenitor cells.

### The evolution of non-ectodermal neurogenesis

Non-ectodermal neurogenesis occurs in several taxa (**Figure 1A**), but it is not well understood to what extend non-ectodermal neurogenesis differs from ectodermal neurogenesis within a given species and whether there are specific similarities in the regulation of non-ectodermal neurogenesis between different taxa. *SoxB* genes have been shown to be involved in ectodermal and non-ectodermal neurogenesis in sea urchin, *Nematostella* and mice, suggesting a general role in neurogenesis (Bylund et al., 2003; Garner et al., 2016; Garriock et al., 2015; Graham et al., 2003; McClay et al., 2018; Richards and Rentzsch, 2014; Richards and Rentzsch, 2015; Tsakiridis et al., 2014; Wei et al., 2011). The identification of *NvPrdm14d* as a gene involved almost exclusively in endodermal neurogenesis in *Nematostella* provides an example for germ-layer specific differences in neurogenesis. While this observation is consistent with other cases of non-ectodermal neurogenesis, such as in the mesodermal pharyngeal neurons of *C. elegans* and the endodermal foregut neurons of the sea urchin (Luo and Horvitz, 2017; Wei et al., 2011), we are not aware of conserved regulators that would be specific for non-ectodermal neurogenesis. The role of *Prdm14* in zebrafish motoneuron development (Liu et al., 2012) shows that its function is not limited to non-ectodermal neurogenesis. However, the role of *Prdm14* in neurogenesis has been poorly described in general and it will be interesting to determine whether this gene is involved in non-ectodermal neurogenesis in other organisms. Differences in the regulation of ectodermal and non-ectodermal neurogenesis can be due to the embryonic origin of the neurons or to the neural cell types that are produced and broader investigations of non-ectodermal neurogenesis in cnidarians and bilaterians are, therefore, required to better understand the evolution of this process. The panel of candidate genes with a role potentially restricted to endodermal neurogenesis in *Nematostella* may serve as a resource for such studies.

### *NvPrdm14d*^+^ neurons as potential motoneurons

The localization of *NvPrdm14d*::GFP^+^ neurons in close vicinity of retractor muscles, their contact with putative post-synaptic sites of the retractor muscles and the expression of *NvIslet* (a direct target of Prdm14 in zebrafish motoneurons), indicate that *NvPrdm14d*^+^ NPCs generate motoneurons in *Nematostella*. The transcriptome analysis of the *NvPrdm14d*::GFP^+^ cells identified additional transcription factors that might be involved in the development of these putative motoneurons, including some with roles in motoneuron development in other organisms [*e.g*. *Tbx20*,(Takeuchi et al., 2005)]. Functional analyses of these transcription factors will likely help identifying conserved and divergent regulators of motoneuron development. How the potential motoneurons are connected to other parts of the nervous system, remains unclear. The retractor muscle functions to retract the head and tentacles into the body column and accordingly the putative motoneurons might connect to the head region via neurites oriented along the oral-aboral axis within the mesenteries, rather than towards the body wall. We can, however, not exclude the possibility that we have missed connections towards the body wall in our sections or that they are established by a population of neurons that is not labelled by either the *NvPrdm14d*::GFP or the *NvElav1*::mOrange reporters.

In conclusion, we identified a population of endoderm-specific, *NvPrdm14d*-expressing neural progenitor cells whose progeny includes potential motoneurons. These observations provide new insights into the regulation of non-ectodermal neurogenesis and open new opportunities for a better understanding of nervous system evolution.

## MATERIALS AND METHODS

### Nematostella vectensis culture

The *Nematostella vectensis* culture is derived from CH2 males and CH6 females initially cultured by (Hand and Uhlinger, 1992). The culture is maintained at 18°C in 1/3 filtered sea water [*Nematostella* medium (NM)], and spawned by a light and temperature shift (18°C to 25°C) for 12 hours as previously described by (Fritzenwanker and Technau, 2002). Fertilized eggs are extracted from their jelly package by incubation in 3% cysteine in NM for 20 minutes with gentle shaking followed by four washes in NM. Embryos are then reared at 21°C and fixed at 24 hours post-fertilization (hpf; early gastrula), 28hpf (mid-gastrula), 30hpf (late gastrula), 48hpf (early planula), 60hpf & 72hpf (mid-planula), 96hpf (late planula), 5 days post-fertilization (dpf; early tentacle bud stage), 6dpf (late tentacle bud stage), 10dpf (primary polyp), or between 2 and 4 months post-fertilization (mpf; growing polyps).

### Cloning of NvPrdm14 paralogs, NvAtonal/neuroD, NvAshD and NvPIT1

The four *NvPrdm14* paralogs were identified by (Vervoort et al., 2015) who established their phylogeny. Sequences were BLASTed to the *N. vectensis* NVE gene models (https://figshare.com/articles/Nematostella_vectensis_transcriptome_and_gene_models_v2_0/807696): *NvPrdm14a* (NVE22869, v1g104327), *NvPrdm14b* (NVE19092, v1g197426), *NvPrdm14c* (NVE9426, v1g61034), *NvPrdm14d* (NVE17327, v1g96522).

The mRNA material of pooled embryos at 24, 48 and 72hpf were extracted using the RNAqueous^TM^ kit (AM1931). The SuperScript^TM^ III First-Strand Synthesis System (Invitrogen, 18080051) was used to generate cDNA from extracted mRNAs to perform PCR amplification. The cDNA of *NvPrdm14* paralogs were cloned into the pGEM^®^-T Easy vector system (Promega, A1360).

*NvAtonal/neuroD* (NVE18219, v1g220027), *NvAshD* (NVE2444, v1g210540) and *NvPIT1* (NVE1681, v1g112929) have been cloned following the same protocol. Digoxigenin-labelled (DIG; Roche 11277073910) and fluorescein-labelled (Fluo; Roche 11685619910) riboprobes were synthetized from these clones with the MEGAscript^TM^ T7 or SP6 kit (Invitrogene, AMB1334/AMB1330).

**Table 1:**
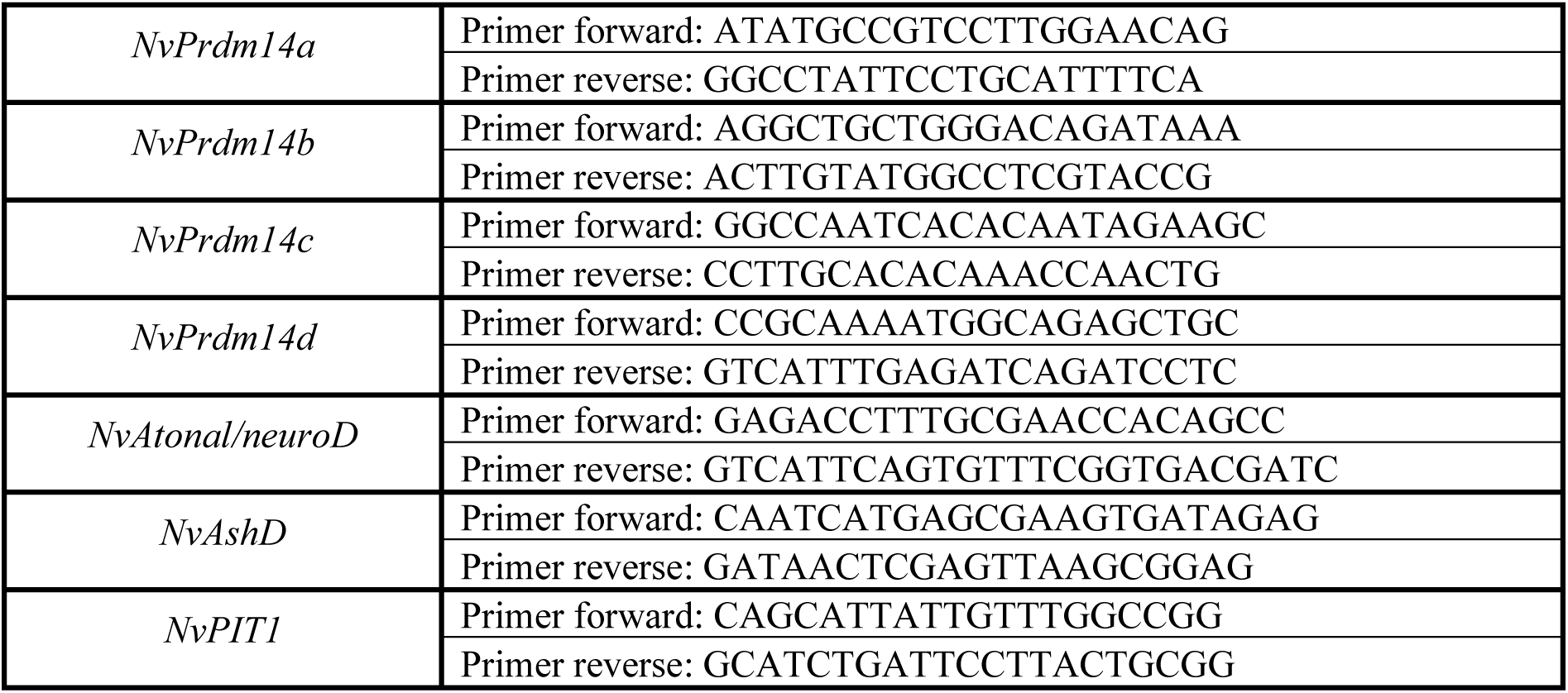
List of primers used for cloning genes.

### Colorimetric (ISH) and fluorescent (FISH) *in situ* hybridization

The following protocols are based on published methods (Richards and Rentzsch, 2014; Tournière et al., 2020) with slight modifications.

Embryos were fixed in 3.7% formaldehyde/0.25% glutaraldehyde/NM for 1 min 30s on ice, then in 3.7% formaldehyde/NM for 1 hour at 4°C. Embryos were washed in PBTw [Phosphate Buffered Saline (PBS)+0.1% Tween20], in PBS, in H_2_O and stored at -20°C in methanol.

For ISH, samples were rehydrated in PBTw, and then incubated in 20 µg/ml proteinase K for 5 min at room temperature (RT) followed by washes in 4 mg/ml glycine/PBTw. They were then washed in 1% triethanolamine in PBTw, followed by the addition of 0.25%, then 0.5% of acetic anhydride. Samples were next washed in PBTw, then refixed in 3.7% formaldehyde/PBTw, followed by washes in PBTw. Pre-hybridization was performed in hybridization buffer (50% formamide, 5X SSC, 1% SDS, 50 µg/ml heparin, 100 µg/ml salmon sperm DNA, 9.25 mM citric acid, 0.1X Tween20) at 60°C overnight (ON). DIG-labelled riboprobes were incubated with the samples at a final concentration of 0.5 ng/µl at 60°C for at least 60 hours. Unbound probes were removed via a series of 60°C washes of post-hybridization buffer (50% formamide, 5XSSC, 1% SDS, 0.1% Tween20, 9.25 mM citric acid) and 2X SSCT (SSC+0,1% Tween20) solutions [75/25, 50/50, 25/75, 0/100 (v/v)], then 0.2X SSCT, 0.1X SSCT. This was followed by washes at RT in PBTw. Samples were then pre-blocked in blocking solution/PBTw [0.5% Block (Roche 11096176001), 50% PBTw, 50% maleic acid buffer (100 mM maleic acid, 150 mM NaCl)] for 5 min at RT, prior to be blocked 2 hours at RT in 1% blocking solution/maleic acid buffer. Samples were then incubated with anti-DIG alkaline phosphatase (Roche, 1:4000)/blocking solution at 4°C ON. Unbound antibodies were removed by 10 x 15 min washes of PBTx/BSA (PBS+0.2% TritonX/0.1% bovine serum albumin); samples were then washed with staining buffer (100 mM Tris pH9.5, 100 mM NaCl, 50 mM MgCl_2_, 0.1% Tween20) before color was developed via addition of 1:100 NBT/BCIP solution (Roche, 11681451001) in staining solution. When the staining reaction was judged to be complete, samples were washed in staining buffer and H_2_O. Samples were cleared via ON incubation with 100% ethanol. Samples are rehydrated in H_2_O, then washed in PBTw and conserved in 87% glycerol at 4%. Samples were imaged on a Nikon Eclipse E800 compound microscope with a Nikon Digital Sight DSU3 camera. Images were cropped on Adobe Photoshop 2020 and some of them were made of assembled focal planes by focus stacking to show a better representation of the complete expression pattern. Figure plates were made on Adobe Illustrator 2020.

For single/double FISH, samples were incubated in 3% hydrogen peroxide/methanol to quench endogenous hydrogen peroxidase activity. Samples were then rehydrated in PBTw and the ISH protocol (see above) was followed from the proteinase K incubation step until the end of the SSCT washes at 60°C. During hybridization, samples were incubated with either DIG or Fluo- labelled riboprobes at a final concentration of 0.5 ng/µl. After the SSCT washes at 60°C, samples were washed in SSCT/PBTw solution at RT [75/25, 50/50, 25/75, 0/100 (v/v)], and then washed in TNTw (0.1 M Tris-HCl pH7.5, 0.15 M NaCl, 0.1% Tween20) and then blocked in TNT/block [0.5% blocking reagent (PerkinElmer FP1012)/TNT] for 1 hour at RT before ON incubation with anti-DIG (1:100, Roche 11207733910) or anti-Fluo (1:250, Roche 11426346910) horseradish peroxidase. Unbound antibodies were removed by 10x15min TNTx (0.1 M Tris-HCl pH7.5, 0.15 M NaCl, 0.2% TritonX) washes, and samples were then incubated in fluorophore tyramide amplification reagent (TSA^TM^ Plus kit, PerkinElmer, NEL748001KT). After the TSA reaction, samples were washed in TNTw and PBTw. For single FISH, samples were incubated with Hoechst 33342 [Thermo Fisher Scientific, 62249 (1:1000)] in PBTx and mounted in ProLong^TM^ Gold antifade reagent with DAPI (Thermo Fisher Scientific, P36935), while for double FISH, samples were then incubated with 0.1 M glycine pH2.0/0,1% Tween20 to inactivate the peroxidase activity of the first antibodies. Samples were then washed in PBTw and TNTw, before being incubated in TNTw/block for 1 hour followed by an ON incubation with anti-DIG or anti-Fluo horseradish peroxidase (Roche). Post-antibody washes and the TSA reactions were repeated as for the first probe; samples were then washed in TNTx, in PBTx, incubated with Hoechst 33342 [Thermo Fisher Scientific, 62249 (1:1000)] in PBTx and mounted in ProLong^TM^ Gold antifade reagent with DAPI (Thermo Fisher Scientific, P36935). Samples were imaged on a Leica SP5 confocal microscope. Images were extracted from stacks with Fiji and adjusted for brightness/contrast and color balance, but any adjustments were applied to the whole image, not parts. Images were cropped on Adobe Photoshop 2020 and figures were made on Adobe Illustrator 2020.

### Cell proliferation assay

Embryos from desired stages were incubated with 100 µM EdU/DMSO in NM for either 30 min, 2 hours or 4 hours at RT and then fixed immediately as described above. Samples were then used for single FISH. After the final washes in PBTx, EdU incorporation was visualized using the Click-iT^TM^ EdU imaging kit with Alexa Fluor^TM^ 647 (Thermo Fisher Scientific, C10424) following the manufacturer’s protocol. Samples were mounted and imaged the same way as for FISH. Images and figures were made the same way as well. The counting of *NvPrdm14d*^+^ and EdU^+^ cells was performed on serial 1 µm sections through the image stack of whole embryos.

### Generation of the *NvPrdm14d::GFP* and *NvFoxA::mOrange2* transgenic lines

The *NvPrdm14d::GFP* transgenic reporter line was generated by meganuclease-mediated transgenesis as described by (Renfer and Technau, 2017).

The genomic coordinates for the ca. 5 kb regulatory region are 226141-231086 on minus strand of scaffold 43 [http://genome.jgi.doe.gov/Nemve1/Nemve1.home.html, accessed in September 2020 (Putnam et al., 2007)]. This region was cloned in front of a codon optimized *gfp* gene with the addition of a membrane tethering CAAX domain to help visualizing the boundaries and morphology of cells expressing the reporter protein. This reporter cassette was flanked by inverted I-Sce1 sites and cloned into the pUC57 vector backbone (GenScript). Wild-type fertilized eggs were injected with a mix containing: plasmid DNA (20 ng/µl), I-Sce1 (1 U/µl, NEB R0694), Dextran Alexa Fluor^TM^ 568 (100 ng/µl) and CutSmart buffer (1X). The mix was incubated at 37°C for at least 15 min, then injection was performed with a FemtoJet^®^ 4i microinjector (Eppendorf). The genomic coordinates for the *FoxA* promoter are 459458 – 465367 on the minus strand of scaffold 58. The promoter fragment was cloned with AscI and PacI into the transgenesis vector described in Renfer et al., 2010 and the line was generated as described above.

### Immunofluorescence (IF)

For immunostainings, embryos were fixed in 3.7% formaldehyde/0.25% glutaraldehyde/NM for 1 min 30s on ice, then in 3.7% formaldehyde/NM for 1 hour at 4°C. Embryos were washed several times in PBTw and in PBTx, and stored at 4°C in PBTx for short periods of time (no more than a week). Embryos older than 72hpf and growing polyps were anesthetized with MgCl_2_ before fixation to prevent muscular contraction. Growing polyps were quickly killed by 30 µl/ml 3.7% fomaldehyde/NM before fixation and fixed in 3.7% formaldehyde/NM ON (instead of 1 hour) at 4°C. Prior to performing IF, the extremity of growing polyp physa was cut to allow a proper staining of mesenteries.

For IF, samples were washed several times in PBTx during 2 hours at RT, blocked in blocking solution [3% BSA/5% Normal Goat Serum (NGS)/PBTx] for 1 hour at RT and incubated with primary antibodies in blocking solution ON at 4°C. Primary antibodies used are: anti-GFP [mouse abcam ab1218 (1:200) or rabbit abcam ab290 (1:200)], and anti-DsRed [rabbit Clonetech 632496 (1:100)] or anti-mCherry [mouse Clonetech 632543 (1:100)]. Samples were then washed extensively in PBTx (5 washes during at least 2 hours 30 min) at RT, blocked in blocking solution for 1 hour at RT and incubated with secondary antibodies in blocking solution ON at 4°C. If phalloidin staining was performed, Alexa Fluor^TM^ 633 phalloidin [Thermo Fisher Scientific, A22284, (1:50)] was incubated at the same time. Samples were then washed extensively in PBTx (5 washes during at least 2 hours 30 min) at RT, incubated with Hoechst 33342 [Thermo Fisher Scientific, 62249 (1:1000)] in PBTx, washed extensively 5 x 10 min in PBTx, and mounted in ProLong^TM^ Gold antifade reagent with DAPI (Thermo Fisher Scientific, P36935). Samples were imaged on a Leica SP5 confocal microscope. Images were extracted from stacks with Fiji and adjusted for brightness/contrast and color balance, but any adjustments were applied to the whole image, not parts. Images were cropped on Adobe Photoshop 2020 and figures were made on Adobe Illustrator 2020.

### Vibratome sectioning

After immunostaining (see above), whole growing polyps were embedded in gelatin/albumin medium [0.4% gelatin type A (Sigma G1890), 27% albumin (Sigma, A3912), 3.7% formaldehyde, in PBS] at RT. Sectioning was performed at RT in PBTx on a Leica VT1000S vibratome. Sections had a thickness of either 20 µm, 50 µm or 100 µm. Sections were stored in PBTx at 4°C and mounted in ProLong^TM^ Gold antifade reagent with DAPI (Thermo Fisher Scientific, P36935).

Sections were imaged on a Leica SP5 or SP6 confocal microscope. Images were extracted from stacks with Fiji and adjusted for brightness/contrast and color balance, but any adjustments were applied to the whole image, not parts. Images were cropped on Adobe Photoshop 2020 and figures were made on Adobe Illustrator 2020.

### Fluorescence-Activated cell sorting (FACS)

Cell were sorted by a fluorescence-activated cell sorter as previously described by (Torres-Méndez et al., 2019).

About a thousand primary polyps positive for *NvPrdm14d*::GFP at 13dpf were dissociated in 0.25% Trypsin (Gibco, 27250018) in Ca- Mg-free NM (CMFNM; 154 mM NaCl, 3.6 mM KCl, 2.4 mM Na_2_SO_4_, 0.7 mM NaHCO_3_) supplemented with 6.6 mM EDTA (pH7.6-7.8) at 37°C for 30 min. Cells were centrifuged at 800g for 10 min, resuspended in ice cold 0.5% BSA in CMFNM (pH7.6-7.8), filtered through a 40 µm filter and stained with Hoechst 33342 (Thermo Fisher Scientific, 62249) at a concentration of 60 µg/ml at RT for 30 min. Samples were then diluted 1:1 with ice cold 0.5% BSA/CMFNM and stained with 60 µg/ml 7-AAD (BD, 559925) for more than 20 min on ice. Cell sorting was performed on a BD FACSAria II with a 100 µm nozzle.

### RNA extraction and sequencing

The RNAseq of *NvPrdm14d*::GFP^+^ cells was performed as previously described in (Torres-Méndez et al., 2019). After FACS, GFP^+^ and GFP^-^ cells were collected in 0.5% BSA in CMFNM (pH7.6-7.8) and centrifuged at 800g and 4°C for 10 min. Most of the liquid was removed and cells were resuspended in TRIzol^TM^ LS reagent (3:1 reagent to sample ratio, Invitrogen 10296028) in 3 independent sorts. Samples were vortexed extensively and incubated at RT for 5 min before being flash frozen and stored at -80°C. The samples were processed using Direct-zol^TM^ RNA MicroPrep columns (Zymo Research, R2060) following the manufacturer’s protocol, including on column DNAse digestion. The RNA quality was assessed using an RNA 6000 Pico kit (Agilent, 5067-1513) on a Bioanalyzer (Agilent G2939A).

Libraries were generated using the NEBNext^®^ Ultra™ II Directional RNA Library Prep kit for Illumina^®^ (NEB #7760L) with 2.5 ng RNA input (1/300 adaptor dilution, 20 PCR cycles). Libraries were sequenced using a 75 bp single end sequencing on a NextSeq^®^ 500 machine (Illumina).

### Bioinformatic analysis of the *NvPrdm14d* transcriptome

The quality control of raw RNAseq-reads was initially assessed with FastQC software v.0.11.8 (Andrews, 2010). Raw reads were then filtered with fastp v.0.20.0 (Chen et al., 2018) using default settings and mapped with STAR aligner v.2.7.3a (Dobin et al., 2013) with default settings to the *Nematostella vectensis* genome (http://genome.jgi.doe.gov/Nemve1/Nemve1.home.html) (Putnam et al., 2007) using (https://figshare.com/articles/Nematostella_vectensis_transcriptome_and_gene_models_v2_0/807696) for NVE gene models. Downstream analysis were performed using R v.4.0.2 (RC, 2020) and Bioconductor packages (https://www.bioconductor.org/). Aligned reads were counted with “summarizedOverlaps” function from the R package “GenomicAlignments” v.1.24.0 (Lawrence et al., 2013) and only genes with at least 10 counts in at least 3 biological replicates for one condition were kept. Differential expression analysis was performed using the package “DESeq2” v.1.28.1 (Love et al., 2014). Overlaps between up- and/or down- regulated genes across different conditions were assessed using the package “GeneOverlaps” v.1.23.0 (Shen and Sinai, 2020). GO terms enrichment was assessed using the package “ClusterProfiler” v.3.16.0 (Yu et al., 2012).

### shRNA-mediated knockdown of *NvPrdm14d*

shRNAs were designed and synthesized based on published methodologies (He et al., 2018; Karabulut et al., 2019) with slight modifications.

Each primer pair was mixed to a final concentration of 20 µM in a total volume of 20 µl, heated to 98°C for 2 min before slow cooling to room temperature. These solutions (5.5 µl each) were then used as template for a 20 µl reaction with the AmpliScribe^TM^ T7 Transcription Kit (Lucigen, AS3107). The reaction product was treated with DNAseI and shRNAs were purified using the RNA Clean and Concentrator^TM^ Kit (Zymo Reasearch, R1012).

Wild-type fertilized eggs were injected with a mix containing: shRNA (900 ng/µl) and Dextran Alexa Fluor^TM^ 488 (100 ng/µl). The mix was injected with a FemtoJet^®^ 4i microinjector (Eppendorf).

**Table 2:**
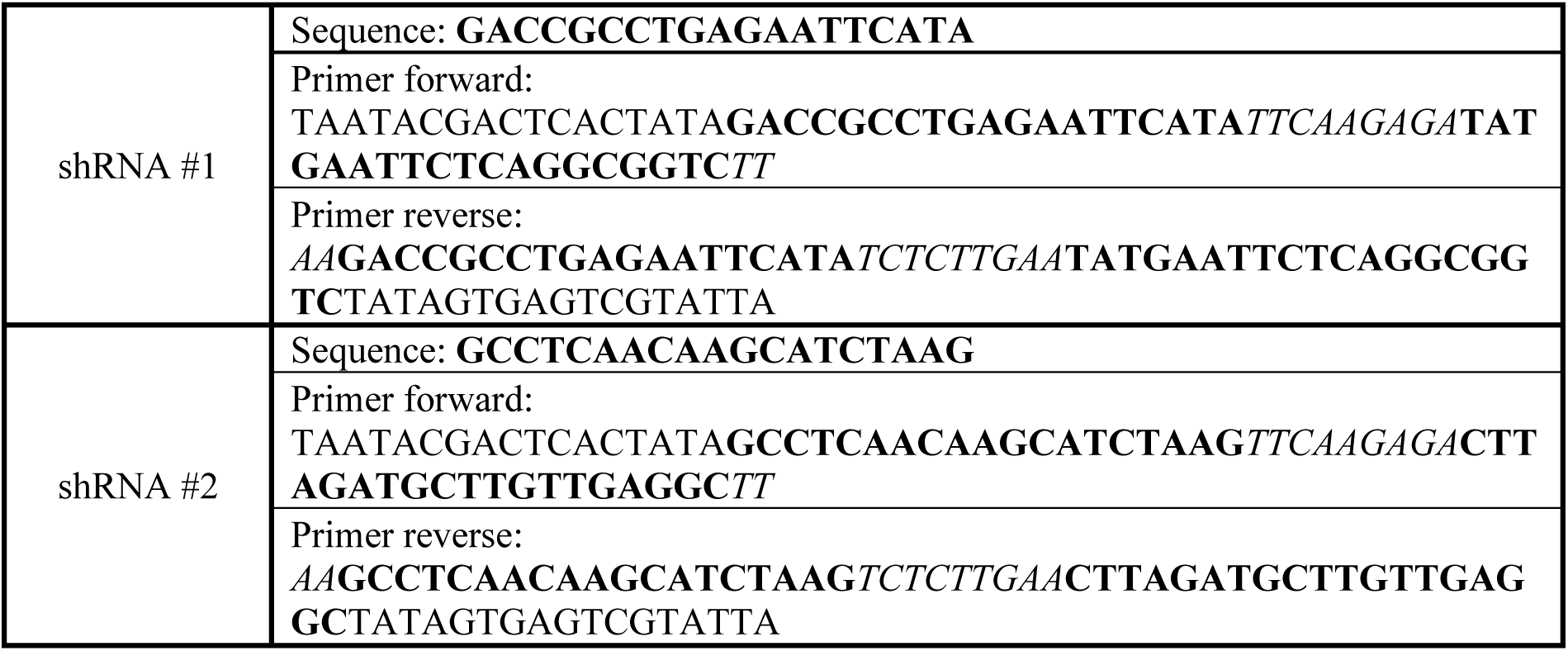
List of primers used to synthetize the different shRNAs targeting *NvPrdm14d*. The sequence of the shRNA is marked in bold. Regular font in primers indicates the T7 promoter sequence and the italic font marks the sequence of the loop found in the final shRNA.

### RNA isolation and qPCR

RNAs from *NvPrdm14d* knockdown were isolated based on a published methodology (Tournière et al., 2022) with slight modifications.

Animals injected with either shRNA #1 or #2 were collected at 48hpf and lysed in 500 µl of TRIzol^TM^ reagent (Invitrogen, 15596026) by vortexing extensively, and incubated at room temperature for 5 min. Chloroform (135 µl) was added and samples were vigorously mixed. The aqueous phase was isolated using MaXtract® High Density tubes (QIAGEN, 129046) according to the manufacturer’s protocol. One volume of 100% ethanol was added to the aqueous phase. This RNA-containing solution was processed using the RNeasy® Mini Kit (QIAGEN, 74104) according to the manufacturer’s instructions, including on column DNAseI digestion using the RNAse-Free DNAse Set (QIAGEN 79254). The SuperScript^TM^ III First- Strand Synthesis System (Invitrogen, 18080051) was used to generate complementary DNA, and it was primed with random hexamers (Custom oligos, Sigma-Aldrich).

qPCRs were performed using the QuantiTect^TM^ SYBR® Green PCR Master mix (QIAGEN, 204143) and ran on a Bio-Rad CFX96 system.

**Table 3:**
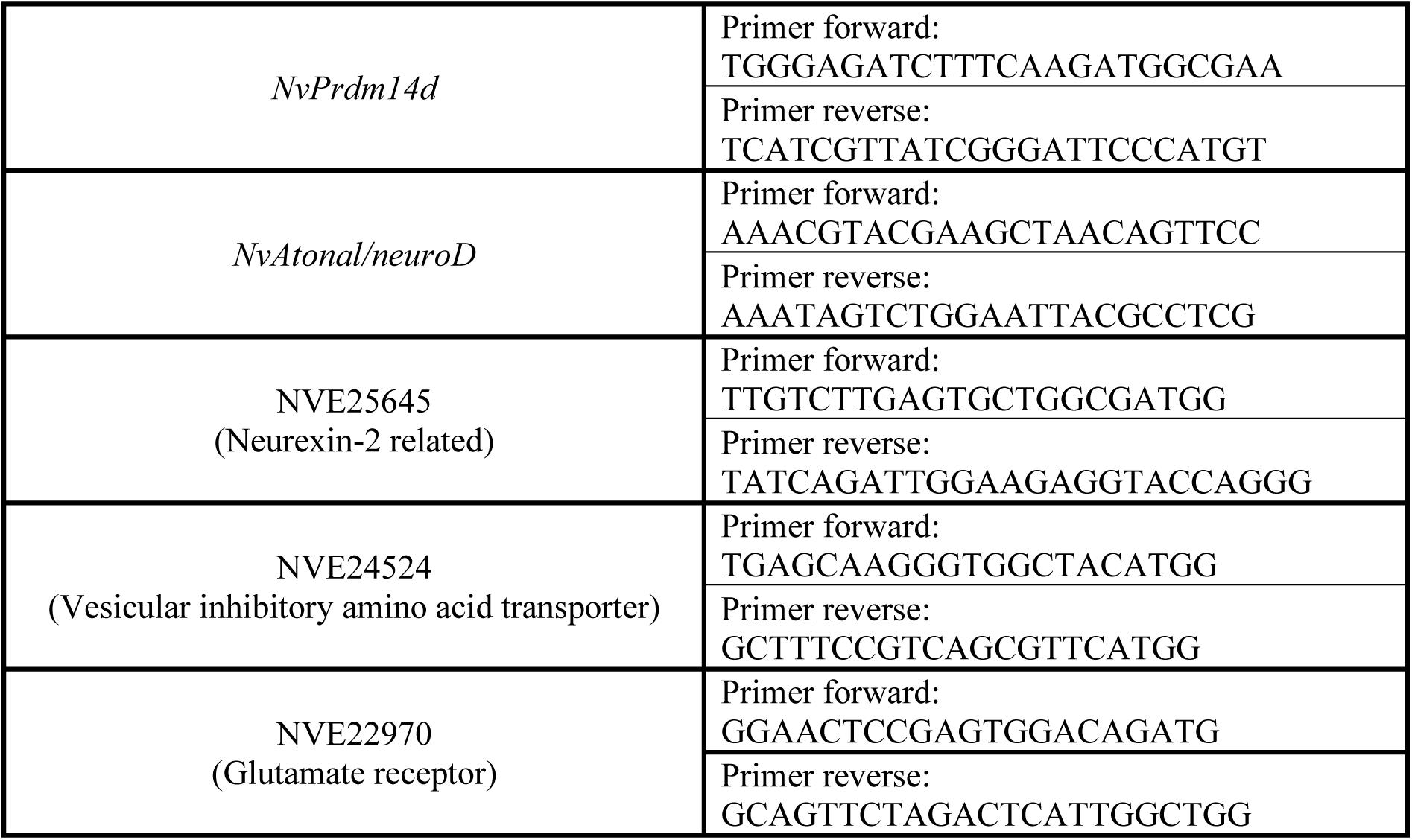
List of primers used for qPCR.

## Acknowledgements

We thank the members of the Rentzsch lab for support and advice, James Gahan for advice and technical help for the generation and analysis of the *NvPrdm14d*^+^ transcriptome, Eilen Myrvold and Lavina Jubek for running the *Nematostella* facility at the Sars Centre, Brith Bergum for performing the cell sorting and Paula Miramón-Puértolas for teaching vibratome sectioning and Anne Kuehnel for experimental help. Cell sorting was performed at the Flow Cytometry Core Facility, Department of Clinical Science, University of Bergen. Sequencing was performed at EMBL GeneCore (Heidelberg). Some of the imaging was performed at the Molecular Imaging Center, Department of Biomedicine, University of Bergen.

## Competing of interests

The authors declare no competing interests.

## Author contributions

Q.I.B.L designed and performed the experimental work, contributed to the conceptualization of the study, analyzed the data, generated the figures, and wrote the manuscript. N.B. performed most of the DFISH and the shRNA injections followed by qPCRs. I.U.K analyzed and visualized the transcriptome data. G.S.R. generated the *NvMyHC1::homer-mCherry* transgenic reporter line and performed the ISH for *NvAshD* and *NvPIT1*. H.B performed some of the DFISH for *NvSoxB(2)*. P.R.H.S generated the *NvFoxA::mOrange2* transgenic reporter line.

F.R. conceptualized and supervised the study and wrote the manuscript.

## Funding

The research reported in the present study was supported by a grant from the University of Bergen and the Research Council of Norway (#251185/F20) and by the Sars Centre core budget to F.R.

## Notes

### Competing Interest Statement

The authors have declared no competing interest.

