## Supplemental figures for "*NvPrdm14d*-expressing neural progenitor cells contribute to non-ectodermal neurogenesis in *Nematostella vectensis*"

### SUPPLEMENTARY INFORMATION

**Figure S1:** A subset of *NvPrdm14d*$^+$ NPCs are dividing.

**Figure S2:** *NvPrdm14d*::GFP$^+$ neurons are closely associated with retractor muscles but not connected to the body wall nerve net through *NvPrdm14d*::GFP$^+$ or *NvElav1*::mOrange$^+$ neuron.

**Figure S3:** A transgenic reporter line for *NvMyHC1*::homer-mCherry highlights putative post-synaptic sites in the retractor muscles.

**Figure S4:** Expression pattern of *NvAtonal/neuroD* by colorimetric *in situ* hybridization.

**Table S1:** Detection of the four *NvPrdm14* genes in available transcriptomic data

**Table S2:** List of genes differentially expressed in *NvPrdm14d*::GFP$^+$ cells.
*See attached Excel file*

**Table S3:** List of genes defining the *NvPrdm14d*$^+$ neuronal metacells.
*See attached Excel file*

**Table S4:** List of upregulated GO terms in the *NvPrdm14d*::GFP$^+$ cells.
*See attached Excel file*

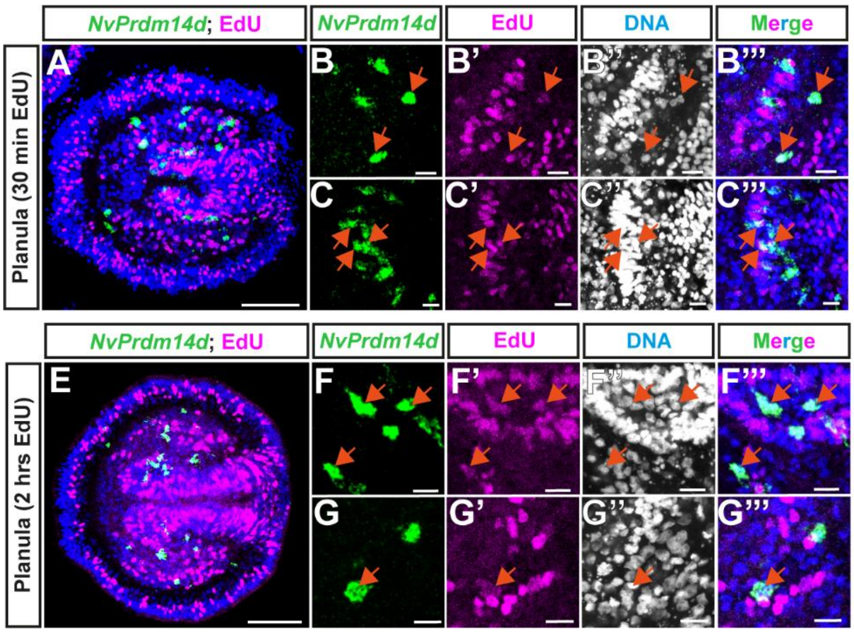


**D**

| | # cells | % |
|---|---|---|
| *NvPrdm14d*+ | 137 | 100 |
| *NvPrdm14d*+ EdU+ | 57 | 41 |

n = 5

**H**

| | # cells | % |
|---|---|---|
| *NvPrdm14d*+ | 243 | 100 |
| *NvPrdm14d*+ EdU+ | 102 | 42 |

n = 5

***Figure S1:*** ***A subset of NvPrdm14d$^{+}$ NPCs are dividing.*** *(**A-C & E-G**) Confocal images of fluorescent in situ hybridization for NvPrdm14d combined with the fluorescent labelling of mitotic cells by nuclear EdU staining, respectively shown in green and magenta. Embryos are incubated in EdU for either 30 min (**A-C**) or 2 hours (**E-G**). (**B-C & F-G**) Enlargements showing that some NvPrdm14d$^{+}$ cells are co-labelled by EdU. (**D & H**) Quantification of the number of NvPrdm14d expressing cells co-labeled by EdU (n = 5). Average numbers of cells per embryo is shown with a ± SD of 25 and 22 for NvPrdm14d$^{+}$ and NvPrdm14d$^{+}$; EdU$^{-}$ cells, respectively (30 min) and with a ± SD of 51 and 31 for NvPrdm14d$^{+}$ and NvPrdm14d$^{+}$; EdU$^{+}$ cells, respectively (2 hrs).*

*Arrows indicate co-labelled cells. Embryos are counterstained for DNA in blue, the oral pole is oriented to the right, scale bars: 50 µm in (**A & E**), 10 µm in (**B-C & F-G**).*

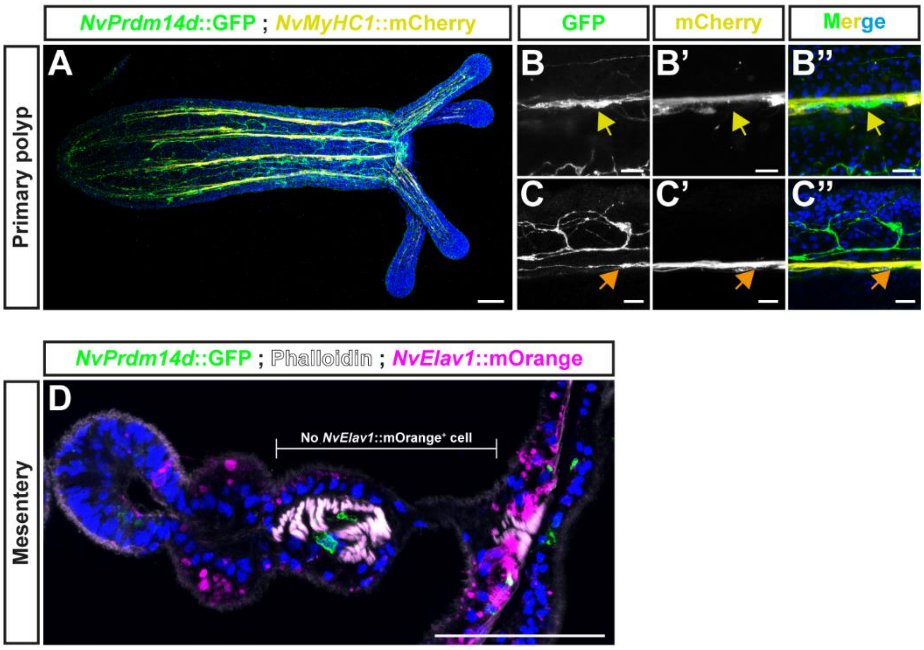


***Figure S2:*** ***NvPrdm14d::GFP+ neurons are closely associated with retractor muscles but not connected to the body wall nerve net through NvPrdm14d::GFP+ nor NvElav1::mOrange+ neuron.*** *(**A-C**) Confocal images of immunofluorescence staining for NvPrdm14d::GFP and NvMyCH1::mCherry, respectively shown in green and yellow. At this stage, neurites expressing NvPrdm14d::GFP run along the retractor muscles. Arrows in (**B-C**) indicate NvPrdm14d::GFP+ neurites associated to retractor muscles. (**D**) Confocal image of immunofluorescence staining for NvPrdm14d::GFP, NvElav1::mOrange and Phalloidin in mesenteries. GFP is shown in green, mOrange in magenta and muscles in white. No NvElav1::mOrange+ neurons is present in the mesenteries between the body wall and the retractor muscles.*

*Samples are counterstained for DNA in blue. The oral pole of the primary polyp is oriented to the right. Scale bars: 50 µm in (**A & D**) and 10 µm in (**B-C**).*

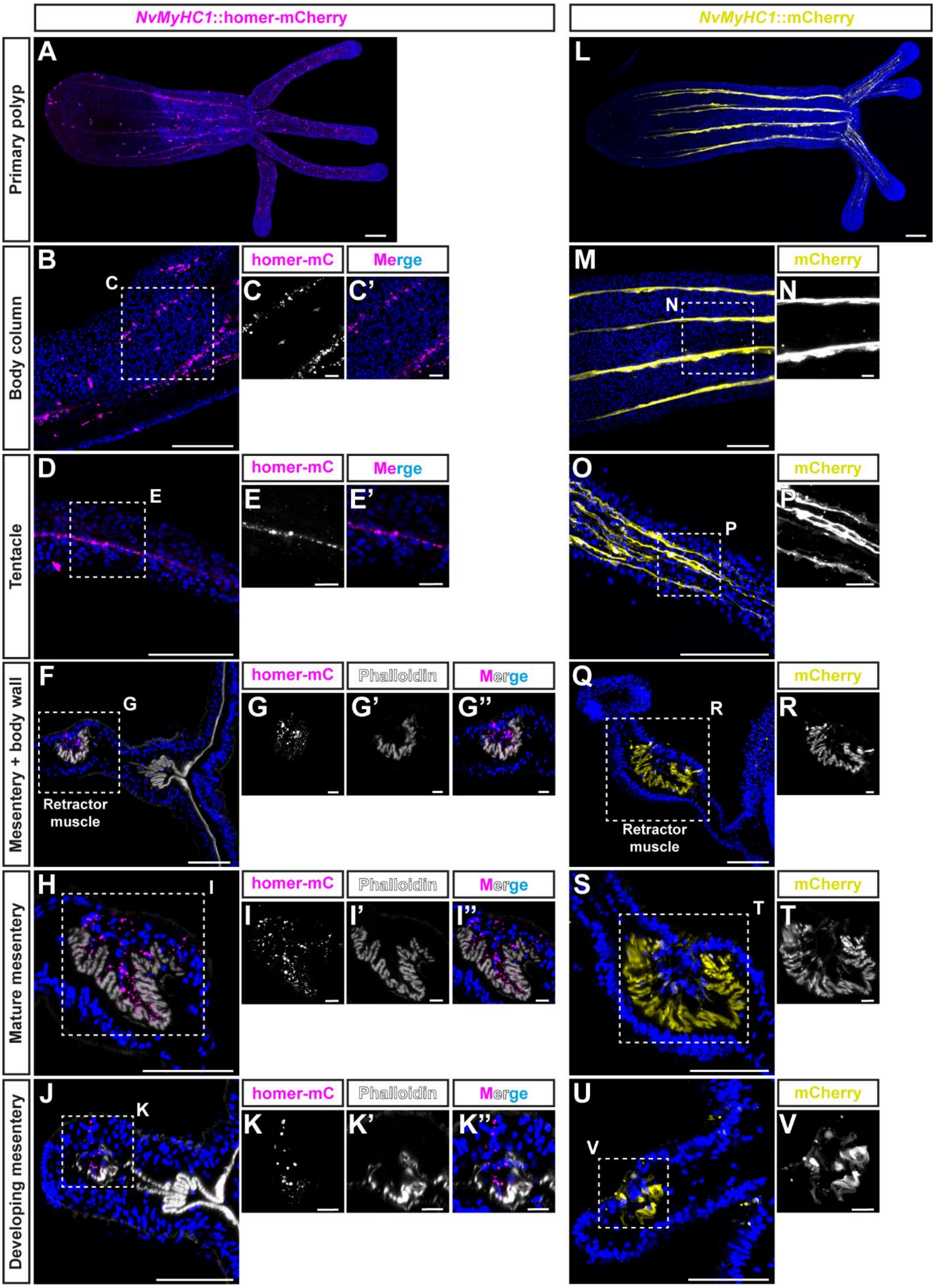


***Figure S3:*** **A transgenic reporter line for NvMyHC1::homer-mCherry highlights the post-synaptic sites in the retractor muscles.** *(**A-V**) Confocal images of immunofluorescence staining for NvMyHC1::homer-mCherry (magenta) and NvMyHC1::mCherry (yellow) in primary polyps (**A & L**), body column (**B-C & M-N**), tentacles (**D-E & O-P**), and different types of mesenteries (**F-K & Q-V**). The Homer-mCherry fusion*

*protein is expressed in puncta along the longitudinal tracts of the primary polyp (**A**), as do retractor muscles shown via the expression of the mCherry protein (**L**). This linear arrangement of Homer-mCherry$^+$ puncta is found in both the body column and the tentacles (**B-E**), a location where mCherry$^+$ retractor muscles are found (**M-P**). Cross-sections reveal that Homer-mCherry$^+$ puncta are exclusively found in mesenteries (**F**), more precisely, associated with retractor muscles (**G-K**). As previously shown by (Renfer et al., 2010), retractor muscles are the only structure labelled by the mCherry protein and they do not exhibit a pattern composed of mCherry$^+$ puncta, whatever the developmental stage of mesenteries (**Q-V**). Therefore, the NvMyHC1::homer-mCherry reporter line is seemingly labelling post-synaptic sites specifically in the retractor muscles.*

*Samples are counterstained for DNA in blue. In (**F-K**), muscles are stained in white by phalloidin. The oral pole of primary polyps is oriented to the right. Scale bars: 50 µm in (**A, B, D, F, H, J, L, M, O, Q, S & U**), 10 µm in (**C, E, G, I, K, N, P, R, T & V**).*

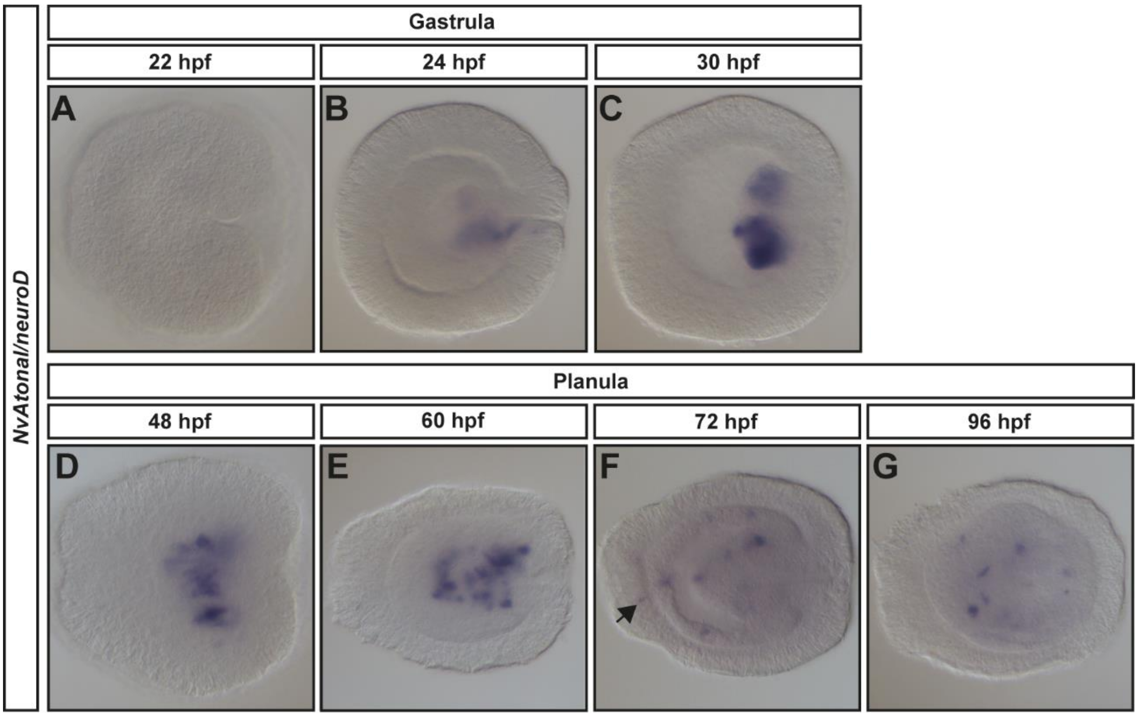


***<u>Figure S4:</u>* Expression pattern of NvAtonal/neuroD by colorimetric in situ hybridization.** *(**A**) In the early gastrula, NvAtonal/neuroD is not expressed. (**B**) In mid-gastrula, NvAtonal/neuroD starts to be expressed in the pharynx. Strong expression in the pharynx persists until mid-planula (**E**). In mid-planula (**F**), NvAtonal/neuroD starts to be expressed in scattered endodermal cells, while the strong expression in the pharynx disappears. Few ectodermal cells express NvAtonal/neuroD (arrow in **F**). From this stage, the expression remains in scattered cells within the pharynx and the endoderm (**F-G**). This expression pattern resembles the one of NvPrdm14d but in fewer cells, suggesting a role for NvAtonal/neuroD in the development of a subset of endodermal NvPrdm14d$^+$ cells/neurons.*
*In all pictures, the oral pole is oriented to the right.*

***<u>Table S1:</u>*** ***Detection of NvPrdm14 genes in available transcriptomic data.*** *The green color indicates where each gene is detected. For the transcriptome of NvPOU4 mutants, it is indicated whether the gene is up- or downregulated. The ID of clusters from the single-cell atlas is indicated when genes are detected. NvElav1$^{+}$ and NvPOU4$^{-/-}$ transcriptomes from (Tournière et al., 2020), NvNCol3$^{+}$ transcriptome from (Gahan et al., 2022), NvPrdm14d$^{+}$ transcriptome from this paper, and single-cell clusters from (Sebé-Pedrós et al., 2018).*

| ***NvPrdm14* genes** | **Transcriptomes** | | | | **Single-cell clusters** | | | |
|---|---|---|---|---|---|---|---|---|
| | ***NvElav1$^{+}$*** | ***NvNCol3$^{+}$*** | ***NvPrdm14d$^{+}$*** | ***NvPOU4$^{-/-}$*** | **Neuronal** | **Cnidocyte** | **Muscle Gastrodermis** | **Gland Secretory** |
| ***NvPrdm14a*** (NVE22869, v1g104327) | | | | **DOWN** | | | | |
| ***NvPrdm14b*** (NVE19092, v1g197426) | | | | | | | | |
| ***NvPrdm14c*** (NVE9426, v1g61034) | | | | | **C34** | | | |
| ***NvPrdm14d*** (NVE17327, v1g96522) | | | | | **C35 & C36** | | | |
